# Functional targeting of Glypican-4 by a conformation-specific single-domain antibody

**DOI:** 10.1101/2025.05.10.653258

**Authors:** Remi Bonjean, Brigitte Kerfelec, Patrick Chames, Rosanna Dono

## Abstract

The heparan sulphate proteoglycan, Glypican-4 (GPC-4), is an integral component of cell surfaces that fulfils key functions as a modulator of cell communication. Over time, human GPC-4 (hGPC4) has gained recognition as a valuable target for enhancing the therapeutic potential of human pluripotent stem cells (hPSCs). hGPC-4 is also a promising diagnostic and therapeutic target for a range of developmental and neurological disorders, as well as cancer. Its involvement in multiple biological processes and its impact on cellular signaling pathways make it a compelling candidate for future research and clinical applications. Here, we report RB1 and RB3 as the first hGPC-4-specific nanobodies, exhibiting high affinity for both recombinant and cell surface-associated hGPC-4 molecules. Notably, the bivalent nanobody Fc-fusion form of RB1, termed RB1-Fc, demonstrates a significant ∼14-fold increase in apparent binding affinity on cells when compared to the monovalent RB1. Furthermore, binding of RB1-Fc to hGPC-4 is dependent on the native conformation of hGPC-4, demonstrating that RB1-Fc is a conformational nanobody. Notably, RB1-Fc neutralizes the activity of GPC-4, as shown by our functional studies in hPSCs. These studies demonstrate the potent efficacy of the lead hGPC4 nanobodies, RB1-Fc and RB3. They also provide a solid rationale for using these nanobodies in the detection and characterization of physiologically and clinically relevant hGPC-4. Additionally, their potential as agents for therapeutic targeting of hGPC-4 opens new avenues for treating disorders associated with dysregulated hGPC-4 activity.

**Highlights:**

- Discovery and generation of nanobodies targeting human Glypican-4.
- RB1-Fc, a bivalent Fc-fusion nanobody, selectively binds to the native human Glypican-4 with high affinity.
- RB1-Fc binding to human Glypican-4 enhances human induced pluripotent stem cells differentiation ability, mimicking human Glypican-4 downregulation
- RB1-Fc is a promissing tool for detecting, characterizing, and therapeutically targeting human Glypican-4 in developmental, neurological, and cancer-related contexts.

**Graphical abstract:** Schematic illustration of this study. Nanobodies (Nbs) specific for native human Glypican-4 (hGPC4) were isolated from a phage-display library generated after immunizing a llama with membrane extracts from hGPC4-transfected HEK 293 cells. Two nanobodies, RB3 and RB1, bind recombinant and cell-expressed hGPC4 with nanomolar affinity. A bivalent RB1-Fc fusion was engineered and showed enhanced binding to endogenous hGPC4 via a conformational epitope. The specificity of the RB1-Fc Nb for hGPC4 is further supported by functional studies in hiPSCs, which demonstrated its blocking activity. Differentiation analyses revealed that RB1-Fc– treated hiPSCs exhibited a significantly enhanced capacity to differentiate into endoderm, closely resembling the phenotype observed in hiPSCs with downregulate hGPC4. These results suggest that RB1-Fc binding functionally inhibits hGPC4, potentially acting as an orthosteric competitor or an allosteric negative modulator.

## INTRODUCTION

Cell signalling serves as a fundamental mechanism for directing cell fate decisions, tissue patterning during embryonic development, and the regulation of organ function and regeneration in adulthood [1, 2]. Over the past decade, research has revealed that a relatively small group of secreted proteins orchestrates many of these complex biological programs [3–6]. Key among them are members of the Fibroblast Growth Factor (FGF), Wnt, Sonic Hedgehog (SHH), and Bone Morphogenetic Protein (BMP) families. These signalling molecules can elicit diverse cellular responses—ranging from fate specification and proliferation to survival, migration, cytoskeletal dynamics, and polarity—highlighting their central role in both development and tissue homeostasis [7]. Mechanistic studies have revealed that the unique signalling properties of secreted signalling proteins are often shaped by their interactions with components of the extracellular matrix [8, 9]. Among these, heparan sulphate proteoglycans (HSPGs)—particularly the glypican family—have emerged as key modulators of these signalling activities, owing to their structural characteristics [9–11]. In mammals, glypicans comprise six members, each consisting of a conserved core protein decorated with two heparan sulfate (HS) glycosaminoglycan chains attached near the C-terminus [9–11]. These proteoglycans are tethered to the cell surface via a glycosylphosphatidylinositol (GPI) anchor but can also be released into the extracellular milieu through proteolytic cleavage, allowing them to influence signalling in both membrane-bound and soluble forms [9–11]. Glypicans have been shown to fulfil three central roles in modulating extracellular signalling. First, they function as co-receptors for key developmental pathways—including Wnt, FGF, SHH, and BMP—by facilitating or hindering ligand-receptor interactions at the cell surface [10, 12]. Second, Glypicans regulate the extracellular distribution of signalling molecules, helping shape morphogen gradients essential for tissue patterning [10, 12, 13]. Third, once released into the extracellular environment—through enzymatic cleavage or shedding—soluble glypicans can modulate ligand availability, influence the spatial range of signalling, and alter the duration of signal activity in neighbouring cells [10, 12, 14]. Consequently, functional studies manipulating Glypican expression—either through loss- or gain-of-function approaches—have underscored their critical roles in development and revealed their involvement in a wide spectrum of human diseases, including genetic syndromes, neurological disorders, and various forms of cancer [15–19].

In recent years, modulating Glypican activity has gained traction as a strategy to fine-tune signaling pathway dynamics under physiological conditions and to mitigate disease progression. For example, Glypican-3 (GPC3) has emerged as a prominent therapeutic target in hepatocellular carcinoma [20, 21], while Glypican-2 (GPC2) blockade is under investigation for its potential in treating neuroblastoma [22, 23]. We have demonstrated that another Glypican, Glypican-4 (GPC4) is an attractive target for modulating stem cell (SC) behavior. We found that reducing GPC4 levels in SCs influences their self-renewal and differentiation capacities [24]. Specifically, we demonstrated that both human induced pluripotent stem cells (hiPSCs) and mouse embryonic stem cells (mESCs) with reduced GPC4 expression transition into a distinct biological state, referred to as the "safe-PSC state". This state is marked by unique biophysical features, enhanced differentiation capacity, and improved therapeutic performance in transplantation-based treatments for Parkinson’s disease [25–28]. Notably, in the context of Parkinson’s disease, hiPSCs with downregulated human GPC4 (hGPC4) protein levels exhibited a significantly greater tendency to differentiate into ventral midbrain dopaminergic neurons (VMDANs) – both in vitro and following transplantation into the brains of rats – demonstrating their potential for more efficient and targeted neural repair [25]. Finally, hiPSCs with downregulated hGPC4 protein levels display markedly lower tumorigenic potential in vivo, indicating that hGPC4 downregulation not only enhances the efficiency of VMDAN differentiation but also significantly improves the safety profile of hiPSC-derived transplants by minimizing the risk of tumor formation after brain engraftment [25].

Beyond stem cell biology, hGPC4 has emerged as a compelling therapeutic target across a spectrum of human diseases. In colorectal cancer, proteasome-mediated ubiquitination of hGPC4 suppresses tumorigenesis by dampening β-catenin/c-MYC signaling pathways [29]. Similarly, in pancreatic cancer, loss of hGPC4 activity enhances the sensitivity of cancer cells to chemotherapy and reduces their stem cell– like characteristics [30]. Moreover, elevated circulating hGPC4 levels have been linked to poor clinical outcomes in glioblastoma, colorectal cancer, and metastatic breast cancer, further underscoring its potential as a prognostic biomarker and therapeutic target in oncology [31, 32]. Elevated serum levels of hGPC4 have been linked to a range of disorders, including central nervous system conditions like Alzheimer’s and Parkinson’s diseases [33, 34], metabolic diseases such as insulin resistance, high body fat content, and non-alcoholic fatty liver disease [35, 36], as well as serving as a novel predictor of all-cause mortality in heart failure patients [15]. Taken together, these findings underscore hGPC4 as a promising therapeutic target for various diseases. Additionally, hGPC4’s association with disease progression positions it as a valuable diagnostic and prognostic marker for disease prediction, patient stratification, and the assessment of therapeutic efficacy.

In current clinical practice, antibodies are widely used for diagnosing and treating various diseases [37]. Beyond traditional antibodies, camelid species (such as llamas) produce heavy chain-only antibodies, which lack light chains [38]. These antibodies’ variable domains, known as VHH or nanobodies (Nbs), retain the antigen-binding specificity of full antibodies and offer several advantages. Nanobodies are significantly smaller (15 kDa compared to the 150–160 kDa of conventional antibodies) and exhibit lower immunogenicity. Furthermore, they possess unique binding properties, high solubility, and exceptional stability, making them a promising alternative to traditional antibody-based therapies [38]. Thanks to their convex-shaped paratope, nanobodies can access and bind to small clefts and cavities [39, 40], as well as conformational epitopes that are typically inaccessible to traditional antibodies [41, 42]. This unique feature enhances their potential as blocking reagents. Single-domain antibodies targeting GPC3 and GPC2 have been successfully isolated from an engineered human VH single-domain phage display library [23, 43]. Here we report the generation of the first two hGPC4-targeting Nbs, namely RB3 and RB1, which specifically bind to recombinant hGPC4 and to hGPC4-transfected cells with nanomolar affinities. We further created a bivalent form of RB1 by fusing it to the Fc domain of human IgG1. This RB1-Fc fusion demonstrated high affinity and specificity for endogenously expressed hGPC4 and recognized a conformational epitope likely located in the core protein of hGPC4. Functional studies revealed that RB1-Fc effectively blocks hGPC4 activity, promoting enhanced endodermal differentiation in hiPSCs. We anticipate that these hGPC4-targeting Nbs could serve as valuable tools to enhance the therapeutic potential of hPSCs without the need for genetic manipulation. Additionally, they may offer therapeutic advantages in treating diseases associated with altered hGPC4 levels.

## Materials and Methods

### Cell Culture and cell transfection

029 hiPSCs[44] and 029 hiPSCS with down regulated hGPC4 [25, 28] were maintained as previously described [25, 28]. HEK 293 (ATCC CRL-1573), Hela (ATCC CRM-CCL- 2), Huh-7 (RRID:CVCL_0336) and SNU-449 (ATCC CRL-2234) were grown as monolayers in Dulbecco’s modified Eagle’s medium (Gibco, ref. 21969035) supplemented with 10% fetal bovine serum (Gibco, ref. A5256801), 1% Penicillin-Streptomycin (Gibco, ref. 15140122) and 1 mM Sodium Pyruvate (Gibco, ref. 11360070) and MKN-45 (RRID:CVCL_0434) in Roswell Park Memorial Institute-1640 medium (Gibco, ref. 11875093) supplemented with 10% fetal bovine serum, 1% Penicillin-Streptomycin and 1 mM Sodium Pyruvate, at 37°C with 5% CO2.

For transfection assay, HEK or Hela cells were seeded the day before transfection and transfected at 70% confluency. Transfection of the plasmids (around 10 µg for 2 x 10^6^ cells in a 10cm tissue culture plate, Corning-Falcon ref. 353003) encoding hGPC4 (Sino Biological, ref HG10090-UT), C-HA-tagged hGPC4 (Sino Biological, ref. HG10090-CY), N-HA-tagged hGPC4 (Sino Biological, ref. HG10090-NY) and mouse Flag-tagged TIGIT (used as irrelevant control molecule; Sino Biological, ref MG50939- NF) was done with Lipofectamine3000 (Invitrogen, ref. L3000015) according to manufacturer’s instruction. Cells were harvested 48 hours after the transfection. Expression of hGPC4 and hTIGIT was analysed by immunocytochemistry using mouse anti-hGPC4 (1:200, Table S1) and mouse anti-FLAG (1:1000, Table S1) antibodies.

### Immunization of llama with hGPC4-enriched cell membranes and Nb library construction

The hGPC4 antigen was prepared from C-HA-tagged hGPC4 transfected HEK293T cells. Two days after transfection, cells were lysed with a native lysis buffer (10 mM Tris, 1 mM EDTA and 5 mM MgCl2) to preserve the conformation of the hGPC4 membrane proteins. Total cell membranes were prepared by ultracentrifugation 80000 g at 4°C for 50 minutes and resuspended in PBS. Llama immunizations were executed in strict accordance with good animal practices, following the EU animal welfare legislation law and were approved by local authorities (French Ministry of Higher Education for Research and Innovation). A llama (lama glama) was injected subcutaneously with membranes prepared from 1.3 × 10^8^ transfected cells on days 0, 9, 18, 28 (Figure S1A and S1B). On day 28 (P4) and day 42 (P5), blood was collected for lymphocyte preparation (Figure S1C).

Library was constructed in the phagemid vector pHEN-phoA-8HisGS as described in Behar et al. 2009 [45]. In brief, total RNA from peripheral blood lymphocytes was reversed transcribed using theTranscriptor Reverse transcriptase (20U/µl, Roche, ref. 03531252103) and the oligo 3’ CH2-2 (5’-GGTACGTGCTGTTGAACTGTTCC-3’). The Nb-encoding sequences were amplified by two successive PCR rounds, digested with SfiI and NotI, and cloned into the SfiI and NotI sites of the phagemid vector pHEN- phoA-8HisGS (pBat379; [45]). P4 and P5 libraries displayed a diversity/complexity of 1.59x10^9^ clones and 4.16x10^8^ clones respectively. ∼99% of transformants in each library harboured a vector with the right insert size (Figure S1D and S1E).

### Phage display and panning method

P4 and P5 libraries were pooled and grown in 50 mL of 2YT medium containing 100 μg/mL ampicillin (2YTA) at 37 °C with shaking at 230 rpm. When bacteria reached OD_600_ between 0.4–0.6, they were infected with helper phage KM13 using a multiplicity of infection of 5×10^9^ pfu/mL for 30 min at 37 °C without shaking. The culture was centrifuged for 15 min at 3000xg, and bacterial pellet was re-suspended in 250 ml of 2YTA with kanamycin (100 μg/mL) for an overnight phage-nanobodies production at 30 °C with shaking. After centrifugation, the supernatant was collected to precipitate the phages with 1/5 (vol/vol) cold solution of 20% PEG8000 and 2.5 mM NaCl 1 hour at 4°C. Phages were spun down and resuspended with 1mL of DPBS 1X. A second round of precipitation was performed for 30 min at 4°C. The phage pellet was resuspended with DPBS 1X and stored at -80 °C with 20% glycerol.

Four rounds of panning were performed by ELISA on Nunc MaxiSorp™ ELISA 96 flat bottom microplates (Nunc/Thermo Fisher Scientific, ref. 442404) pre-coated with the recombinant hGPC4-Fc chimera protein (4.4 µg of hGPC4-Fc in DPBS 1X (0.4 µg per well); R&D Systems, ref. 9195-GP-050) overnight at 4°C with shaking. After 3 washes with PBS/0.1% Tween-20, the plates were treated with blocking buffer [3%(wt/vol) milk in DPBS 1X] at room temperature for 1 h with shaking. After two washes with DPBS 1X containing 0.1% Tween-20 and two washes with DPBS 1X, the Fc epitopes were masked as previously described [46] with human anti Fc specifc Nbs (0.2 mg/ml) in blocking buffer at room temperature for 1 h with shaking. The phage library, resuspended in 1mL blocking buffer, was incubated for 90 min at room temperature with the immobilized hGPC4 antigen. After nine washes with DPBS 1X containing 0.1% Tween-20 and three washes with DPBS 1X, bound phages were eluted by treatment with 1 mL of Trypsin (Sigma-Aldrich, ref. T1426) at 1 mg/mL in DPBS 1X for 30 min at room temperature. Eluted phages-Nb were amplified by infection of E. Coli TG1 and produced as described above, for the other rounds of panning. A Fc binders depletion step on Maxisorp plates pre-coated with hIgG1 (11 ug/ml) overnight at 4°C with shaking was added before the third and fourth panning to maximize the selection of hGPC4 binders. Finally, 180 individual Escherichia coli TG1 colonies randomly picked from the selection outputs were grown overnight at 37°C in 2YTAG for the screening step. The production of soluble nanobodies was induced by the addition of 0.1 mM IPTG (Euromedex, ref. EU0008-C) and overnight growth at 30°C. Nb-containing supernatants were harvested and tested by ELISA on Maxisorp plates pre-coated with hGPC4-Fc protein as described below. A threshold absorbance (OD450/405 nm) of 2**-** fold higher absorbance of an irrelevant nanobody was used for the identification of positive hits. The genes of the positive hits were sequenced to characterize Nb diversity (GENEWIZ).

### Production and purification of recombinant Nbs

A representative Nb from each identified clusters was selected, produced after transformation of E. coli BL21 DE3 strain by the selected phagemids and purified as previously published [45]. Briefly, after lysis of bacteria using Bugbuster buffer (Novagen, ref 70584) supplemented with 20 µg/ml Lysozyme (Eurobio, ref. GEXLYS00-6Z) and 25 U/µl Benzonase (Millipore, ref. 70746), Nbs were purified by affinity chromatography on TALON superflow™ cobalt resin (GE Healthcare, 28-9575- 02) equilibrated with DPBS 1X. The resin was washed with 10 column volumes of DPBS 1X, 300 mM NaCl then with 10 column volumes of DPBS 1X. Nanobodies were eluted with DPBS 1X, 150 mM imidazole, desalted on PD-10 and concentrated if necessary (Vivaspin, Sartorius, ref. VS0611). The protein concentration was determined spectrophotometrically (Direct Detect®) and Nbs purity was evaluated by SDS-PAGE and Western blot.

### Generation of the Nb-Fc construct, production and purification

The Nb-Fc fusion was constructed by cloning the Nb coding sequence amplified by PCR into the AgeI/BstEII pre-digested pHLSec- Fc-6His plasmid (Addgene ref. 99846) upstream the Fc sequence using the NEB HiFi kit according to the manufacturer’s instruction (New England Biolabs, ref E2621). Nbs coding sequences were amplified from the phagemids using PCR Phusion (Thermofisher, ref. F530S) and the oligos 5’HLSecVHH: (5’-GGTTGCGTAGCTGAAACCGGTGAGGTGCAGCTGGTG-3’) and 3’VHHEndH (5’-CGGTGGGCATGTGTGAGTTTTGTCTGAGGAGACGGTGACCTG-3’). The Nb-Fc plasmids were then transfected in Expi-293F cells (Gibco, ref. A14635) following manufacturer’s protocol to produce the Nb-Fc proteins. Four days after transfection, the cell supernatant was collected, centrifuged, dialyzed against 1x DPBS (Spectrumlabs - 12-14 kDa cutoff, ref. 132700) and purified on protein A column (GE Healthcare, ref. 17-0403-01) on ÄKTA go™ followed by buffer exchange on PD-10 desalting column (Cytiva, ref. 17085101) before storage in PBS 1X.

### SDS-PAGE and Western Blot Analysis of Nb Production

SDS-PAGE was performed on a 4-20% Mini PROTEAN® TGX Stain-Free™ Protein gel (BioRad) under reducing and non-reducing conditions. The gel was imaged with ChemiDoc Touch imaging system (Biorad). Western blot was performed on nitrocellulose membrane using a trans-Blot Turbo Transfer System (BioRad). After membrane saturation for 1h at room temperature in DPBS 1X supplemented with 5%(wt/vol) BSA, membranes were incubated with anti-6His peroxidase conjugated antibodies (Table S1). Detection was performed with HRP-conjugated anti-HIS mouse antibody (Table S1) and substrate solution (DAB (3,3′-diaminobenzidine tetrahydrochloride) and H2O2). Precision Plus Protein™ unstained, Prestained Standards (BioRad) and PageRuler Plus 10-250KDa (Thermofisher) were used for SDS-PAGE and Western blot respectively.

### ELISA

ELISA was performed on Nunc MaxiSorp™ ELISA 96 flat bottom microplates (Nunc/Thermo Fisher Scientific, ref. 442404) pre-coated overnight at 4°C with either 4 µg/ml of recombinant hGPC4-Fc chimera protein (R&D Systems, ref. 9195-GP-050), or irrelevant proteins (hErbB2-Fc (Sino Biological, ref 10004-H02H) or human IgG). All ELISA steps were subsequently done at room temperature. Saturation was performed with blocking buffer [3% (wt/vol) milk in DPBS 1X] for 1h with shaking, followed by 3 washes with DPBS 1X/0.1% Tween-20. For the screening step, 70 μL of supernatant from TG1 culture was mixed with 30 µl of blocking buffer and added to the pre-coated plate for 1 hour. After incubation, the wells were washed 3 times with DPBS 1X/0.1% Tween-20 and the bound Nbs were detected by the addition of an HRP-conjugated anti-6His mouse antibodies (Table S1). Peroxydase activity was measured at 450 nm after addition of TMB (SeraCare, ref 5120-0077) and HCL 1N stop solution using a Tecan Infinite® M1000 plate reader. To measure the Nbs or Nbs-Fc affinities, serial dilutions of purified Nbs or Nbs-Fc in 3%(w/v) milk/DPBS 1X were added to the recombinant hGPC4-Fc pre-coated Maxisorp plates and incubated for 1 h. The plates were then processed as described above. The coating of hGPC4-Fc protein was controlled by incubation with mouse anti-GPC4 antibodies (Table S1) and revealed with peroxidase-conjugated goat anti-mouse antibodies (Table S1). The coating of irrelevant proteins (herbB2-Fc or hIgG proteins) was revealed with peroxidase-conjugated goat anti-Human antibodies (Table S1).

### Flow cytometry

For flow cytometry, growing cells were washed and harvested with Accumax (Millipore, ref. SCR006). As HeLa cells do not express detectable levels of hGPC4, we used hGPC4-transfected Hela cells (hGPC4-HeLa), whereas non-transfected or TIGIT- transfected HeLa cells (TIGIT-HeLa) were used as negative controls. Following centrifugation, cells were resuspended in ice-cold DPBS1X / 5%(wt/vol) BSA and plated into 96 well-plate at 1.5-3x10^5^ cells per well (Nunc/Thermo Fisher Scientific, ref. 24950). Cells were incubated with various amounts of Nbs or Nbs-Fc for 1 hour at 4°C and then washed twice with DPBS1X/ 5% BSA to remove non specific binding. When testing affinity of Nbs, cells were incubated first with anti-6His primary antibodies (Table S1) for 45 min with shaking and then, following two washes with DPBS1X/ 5%BSA, with Alexa647conjugated goat anti-mouse secondary antibody (Table S1) for 30 min. When testing affinity of Nbs-Fc, cells were incubated with Phycoerythrinconjugated goat anti-human IgG secondary antibodies (Table S1) for 30 min. For each experiment, at least 10,000 cells were analyzed and the fluorescence associated with the live singlet cells was measured using a MACS Quant cytometer (Miltenyi).

### Immunoprecipitation assays

HEK 293 cells transfected and cultured in 10 cm tissue culture plates (with 2 x 10⁶ cells seeded one day prior to transfection) were lysed using 1 mL of lysis buffer per plate. Lysis was performed using either a non-denaturing buffer (referred as non-denaturing buffer 1: 20 mM Tris-HCl, pH 7.5; 150 mM NaCl; 1 mM EDTA; 1 mM EGTA; 1% NP-40) or a buffer supplemented with the ionic detergent, sodium deoxycholate (referred as non-denaturing buffer 2: 20 mM Tris-HCl, pH 7.5; 150 mM NaCl; 1 mM EDTA; 1 mM EGTA; 1% NP-40; 1% sodium deoxycholate), both in the presence of protease inhibitors. These two different non-denaturing lysis buffers were used to test the effects of detergents on the interaction between RB1-Fc and hGPC4. Protein denaturation was achieved by lysing cells in an SDS containing buffer (1%SDS, 5 mM EDTA, 10mM DTT and protease inhibitors). Cells were resuspended in this denaturing lysis buffer, mixed well by vigorously vortexing and heated at 95°C for 5 min to denature proteins. The DNA was fragmented by passing the lysed suspension 5–10 times through a needle attached to a 1 mL syringe. Following centrifugation to remove insoluble material, protein concentration was determined using the Bradford assay (Bio-Rad, Cat. No. 5000006) or, alternatively, by Coomassie Blue staining (Bio-Rad, ref. 1610803). Cell lysates (1.5 mg total protein) were incubated with the indicated antibodies (20 µg) in 500 µL to 1 mL of non-denaturing buffer (volume adjusted based on protein concentration) overnight at 4 °C under gentle rotation. For proteins extracted using a denaturing lysis buffer containing SDS, lysates were diluted 10-fold in non-denaturing buffer prior to antibody incubation to reduce SDS concentration. Packed protein A-Agarose beads (30µl) (TermoFischer Scientific, ref. 20333) equilibrated with non-denaturing buffer were added to the antibody-lysate sample for 2 h at 4°C under rotation. Beads were spun down and washed with non-denaturing buffer five times. The immune complexes were released from the beads by heating at 100°C for 5 min in 100 µl of 6X loading buffer (375 mM Tris-HCl pH 6.8, 6% SDS, 4.8% Glycerol, 9% 2-Mercaptoethanol, 0.03% Bromophenol blue). Western blot analysis was performed on 40 µl of immune complexes under reducing conditions. Total cell lysate input (60 µg) was included as control. Proteins were resolved on 10% Anderson gels (http://www.pangloss.com/seidel/Protocols/proteingel.html) under nonreducing conditions, and transferred to PVDF membranes. After blocking with 5% milk/ PBS/, 0.1% Triton X100, the membranes were incubated overnight at 4°C with the rat anti-HA antibody (Table S1). The membranes were then washed, incubated with HRP-conjugated anti-rat IgG (Table S1) at room temperature for 1hr, and peroxidase activity was visualized with the ECL Plus Kit (Amersham, ref. 11517371).

### Immunofluorescence analyses on cultured cells

Cells grown on glass coverslips or labtek (Nunc, ref. C7182) were fixed for 10 min with 4% PFA in DPBS1X. All steps were done at room temperature unless otherwise specified. After three washes with PBS, fixed cells were either incubated 20 min with 0.1% TritonX-100 (for better staining) in DPBS1X for extra-cellular staining or permeabilized with 0.3% TritonX-100 in DPBS1X for intra-cellular staining. Following 1 hour blocking in 3% BSA (Sigma, ref. A9647), 2% Donkey serum (Abcam, ref. ab7475), 0.3% or 0,1% TritonX-100, in DPBS1X, cells were incubated with primary antibody overnight at 4°C in the same blocking solution. After three washes with DPBS1X / 0.3% or 0.1% TritonX-100, cells were incubated with secondary antibodies (Table S1 + DAPI 5 µg/ml; Roche, ref 10236276001) for 1 hour protected from light. Finally, after three washes with DPBS1X / 0.3% or 0,1% TritonX-100 and two with DPBS1X, coverslips or Labtek were mounted using ProLong™ Gold Antifade Mountant. The images were captured on a Zeiss AxioImager APO Z1 microscope. For immunofluorescence experiments using RB1-Fc, the nanobody was applied at concentrations ranging from 8 to 25 nM.

### hGPC4 quantitative PCR analysis

RNA isolation and cDNA synthesis were performed as previously described[25]. For *hGPC4* quantitative PCR, 2.7 ng of cDNA was amplified using SYBR Green qPCR SuperMix (Thermofisher, ref. 11761500) and 0.1 µM of forward and reverse primers. Levels of all transcripts (Ct) were normalized to those of the housekeeping gene GAPDH (ΔCt) and subsequently to the ΔCt of the A549 cell lines that do not express *hGPC4* (ΔΔCt). Results were reported as relative quantities (RQ = 2^-ΔΔCt). Oligos used for *hGPC4* were: FW 5’-3’ GTCAGCGAACAGTGCAATCAT and REW 5’-3’ ACATTTCCCACCACGTAGTAAC. Oligos used for *GAPDH* were: FW 5’-3’ GTCTCCTCTGACTTCAACAGCG and REW 5’-3’ ACCACCCTGTTGCTGTAGCCAA.

### Cell viability assays

For survival assays, hiPSCs cells were seeded in 96 well-plates (Corning, ref.353377) at 1 x 10^4^ cells per well. Two days after, the medium was changed and replaced with fresh medium containing increasing amount of RB1-Fc. At day 3, cell viability was revealed by addition of CellTiter-Glo reagent (Promega, ref G7570) in cell supernatant. After 20 min of incubation at room temperature (light protected), the luminescence activity was analyzed with a luminometer microplate reader (Berthold). Cell survival was normalized to cells treated with vehicle (PBS).

### Analysis of Nbs-Fc blocking properties: endoderm differentiation in hiPSCs

In vitro differentiation of hiPSCs into definitive endoderm (DE) progenitors was performed according to Legier et al., 2023 [28]. Briefly, hiPSCs were seeded on Matrigel-coated plates or coverslips at a density of 1.1 x 10^5^ cells/cm² in mTeSR1 medium [28] supplemented with 10 μM of ROCK inhibitor Y-27632 (Ri; Tocris, ref. 1254). To induce DE differentiation, hiPSCs were washed twice with RPMI (Invitrogen, ref. 21875034) to remove self-renewal growth factors and treated for 1 day with ACTIVIN A (R&D, ref. 338-AC, 100 ng/mL) in DE medium (RPMI 1640, 1% Penicillin/Streptomycin, 1% L-Glutamine (Gibco, ref. 25-030-081). The day after differentiating, cultures were treated with ACTIVIN A in DE medium supplemented with 0,2% of FBS (Hyclone, ref. SH30071.02E). The hiPSCs were exposed to increasing concentrations of the Nbs RB1-Fc during DE differentiation (50 and 500nM). Two treatment conditions were applied to hiPSCs. In the first, RB1-Fc was added at the time of seeding (day 0); in the second, treatment with RB1-Fc was initiated on the first day of DE differentiation (day 1). In both cases, cells were exposed to RB1-Fc through all differentiation procedure (3 days). The medium was changed daily to keep a constant level of RB1-Fc. Then, cells were fixed and analyzed by immunocytochemistry for the presence of the SOX17 protein using SOX17 antibodies (Table S1).

### Image analysis

To quantify the percentage of SOX17-positive cells over DAPI (used to define nuclear areas), the threshold of each single channel picture was first set using the Zeiss software ZEN (version 3.4). Images were converted to grayscale pixels. The percentage of pixels that were highlighted by the threshold (corresponding to SOX17- and DAPI-positive positive) was determined as area fraction using the FIJI software[47]. Values were reported as ratio of the area fraction of SOX17-positive cells over DAPI area fraction.

### Statistical analysis

Statistical analyses were performed using the most adapted test for each study (e.g. One way ANOVA) using the GraphPad Prism version 8 software. Nb-binding curves were plotted using nonlinear least square fit. Kd values were calculated using the one site binding model reported in the Prism version 8 software. Data are presented as mean with error bars representing standard deviations unless stated otherwise. Statistic values were reported as: ns=not significant, * for p < 0,05, ** for p < 0,01, *** for p < 0,001.

## RESULTS

### Generation of hGPC4-targeting Nbs

HGPC4 nanobodies specific for the native conformation have been isolated from phage-display libraries generated after immunization of one lama with membrane extracts from hGPC4-transfected HEK 293 cells (Figure S1). Phage particles were generated and subjected to panning on a recombinant chimeric hGPC4-Fc fusion. After four consecutive rounds of panning including a depletion step on human IgG1 for the last two rounds, 360 individual clones were isolated and screened by ELISA on recombinant hGPC4-Fc. Based on the OD threshold, 16 potential hGPC4 binders were selected (Figure 1A and Figure S2A). Sequencing of the 16 clones identified 4 different sequences named RB1, RB2, RB3 and vRB3, the last two differing by only one mutation (Figure 1B). The RB1 sequence was identified in 3 independent clones, RB2 in 2 clones, RB3 in 5 clones, and vRB3 in 6 clones.

**Figure 1.**
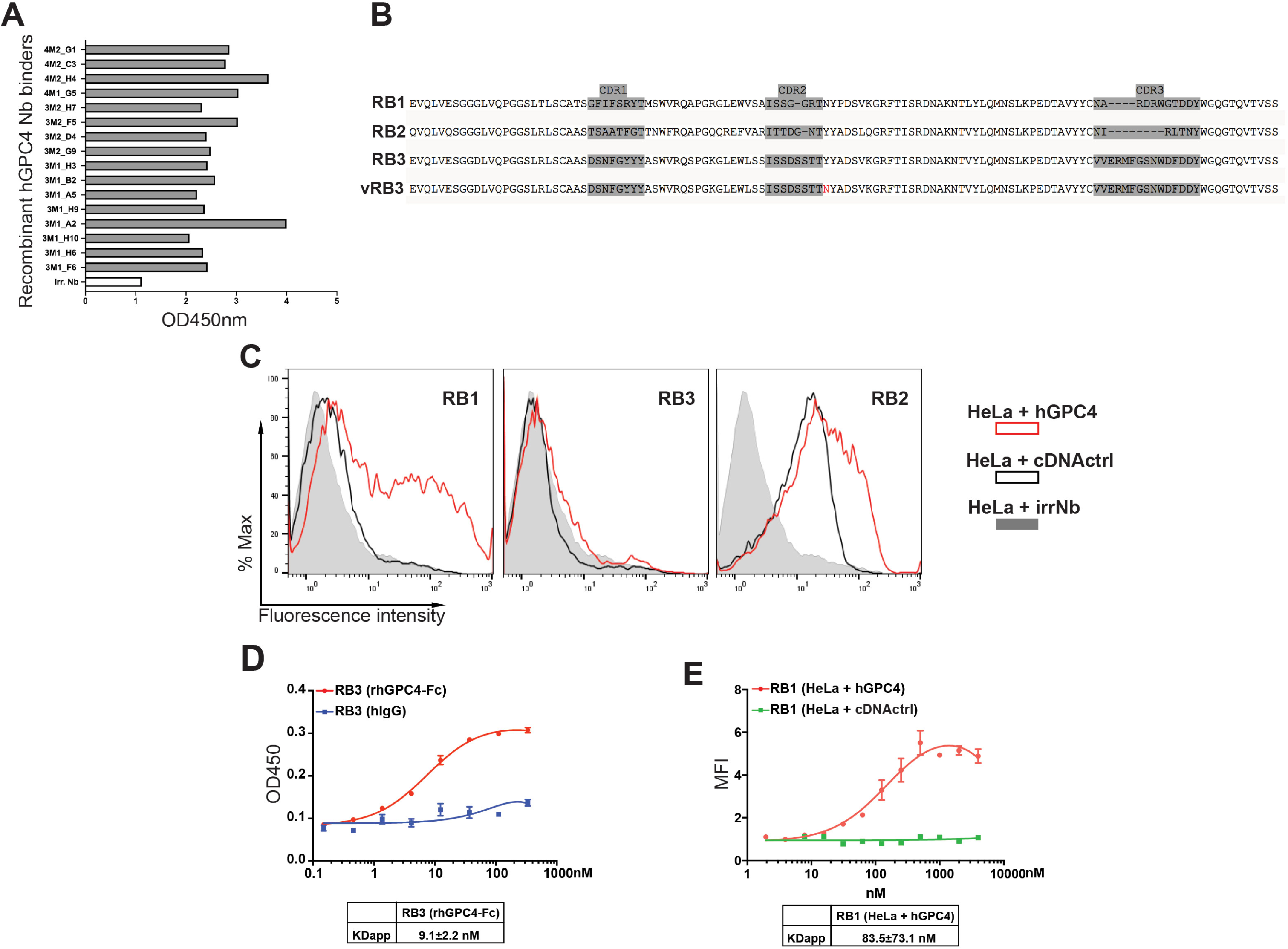
Identification of hGPC4-specific single-domain antibodies by phage display. (**A**) ELISA of the sixteen potential hGPC4 binders selected on recombinant hGPC4-Fc protein. (**B**) Amino acid sequence of the four different Nbs identified. Sequences are numbered according to the Kabat numbering scheme. The complementary determining regions (CDR1, CDR2 and CDR3) were assigned according to the AbM definitions [54] and are shown with a grey background. Based on their amino acid sequences, the Nbs were categorized into four distinct groups: RB1, RB2, RB3, and vRB3. Notably, RB3 and vRB3 are derived from related B-cell clones, differing only by a single amino acid substitution – tyrosine (Y) to asparagine (N) – at the first residue of the framework region 3 (FR3). (**C**) Flow cytometry analysis of binding properties of one representative Nb belonging to the RB1, RB2 and RB3 classes to HeLa cells transfected with hGPC4 expression vectors (red curve), and to HeLa cells transfected with a non-specific plasmid (black curve). Shaded grey areas are the irrelevant Nb (irrNb) signal on the hGPC4 transfected HeLa cells. (**D**) Binding of RB3 to recombinant hGPC4-Fc protein (red curve) and to hIgGs (blue curve) examined by ELISA. RB3 was added at increasing concentrations. Apparent K_D_ were calculated using the GraphPad Prism version 8 software. Data are presented as mean±SEM of n=2. Before pooling, data were normalized by the values of the negative control obtained with a an irrNb. (**E**) Binding of RB1 to HeLa cells expressing either hGPC4 (red curve) or an irrelevant protein (cDNA ctrl; green curve) measured by flow cytometry. Cells were incubated with increasing concentrations of RB1. Apparent K_D_ were calculated using the GraphPad Prism version 8 software. Data are presented as mean±SEM of n=3 biological replicates. Before pooling, data were normalized by the values of the negative control obtained with a an irrNb.

The capacity of the nanobodies (Nbs) to recognize the native conformation of hGPC4 was evaluated by flow cytometry using hGPC4-expressing HeLa cells. Specificity was determined by comparing fluorescence signals between hGPC4-HeLa and control TIGIT-HeLa cells. RB1 exhibited specific binding exclusively to hGPC4-HeLa cells (Figure 1C, Figure S2B). In contrast, RB2 bound both hGPC4- and control-transfected HeLa cells and was thus excluded from further analysis (Figure 1C, Figures S2B and S2C). RB3 and its variant, vRB3, failed to recognize membrane-bound hGPC4 (Figure 1C, Figures S2B and S2D).

Complementary ELISA-based binding assays using recombinant hGPC4-Fc confirmed that RB1 and RB3, but not RB2, bound the recombinant protein (Figure S2E, Figure 1D).

Taken together, these results show that RB1 and RB3 are hGPC4 specific binders: RB1 recognized both recombinant and cell-expressing hGPC4 protein, whereas RB3 similar to vRB3 binds only to recombinant hGPC4 protein.

### Analysis of hGPC4 Nbs

To characterize further the binding properties of RB1 and RB3, we determined their apparent binding affinity for hGPC4. Affinity measurements were performed using serial Nb dilutions of RB3 and RB1 either by ELISA on recombinant hGPC4-Fc or by flow cytometry on hGPCA-transfected HeLa cells, respectively. Titration curves analysis from independent experiments revealed that RB3 bound recombinant hGPC4 with an apparent equilibrium dissociation constant of ∼9.1+/-2.2nM and RB1 bound to cell-expressed hGPC4 with an apparent K_D_ of 84±73.1nM (Figure 1D and 1E). The interaction of RB1 with native hGPC4 expressed by cells suggests potential blocking properties. Given this possibility, RB1 was selected for further analysis.

### Dimerization of the RB1 Nbs enhances its hGPC4-binding ability

The conversion of Nb to IgG format offers several advantages such as increased affinity through avidity, stability in biological fluids, increased half-life and Fc-mediated functions. We generated a human RB1-Fc fusion protein by fusing RB1 in frame with the hinge, CH2 and CH3 domains of human IgG1 (Figure 2A and Fig S3A) resulting in a bivalent 80 kDa RB1-Fc protein, as compared to the 15 kDa size of the monovalent RB1 Nb (Figure 2A and Figure S3B).

**Figure 2.**
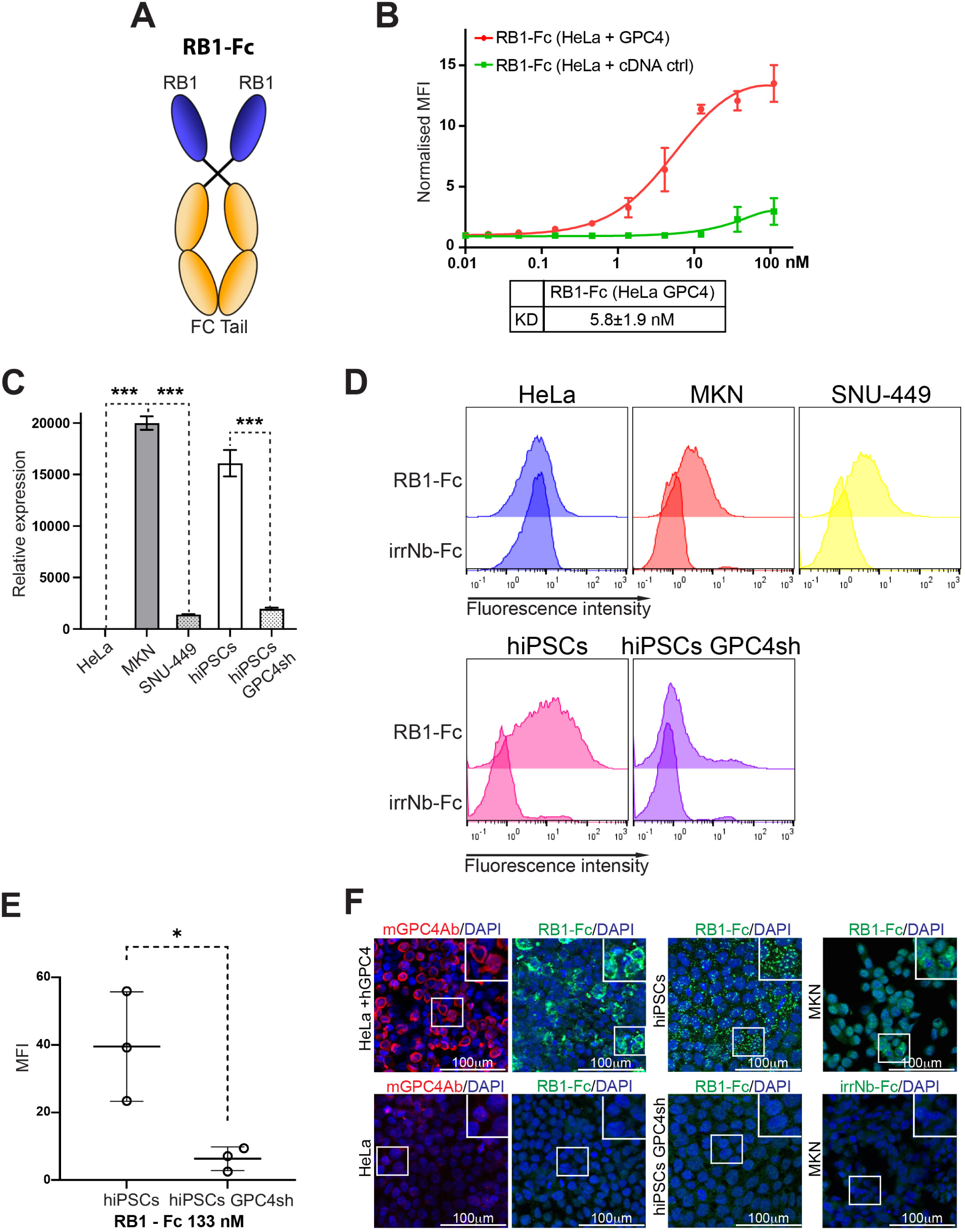
Binding properties of the bivalent RB1-Fc Nb. (**A**) Schematic representation of the bivalent RB1-FcNb generated by genetically cloning the VHH domain of RB1 in frame with the Fc domain of a human IgG1. (**B**) FACS analysis of cell binding on HeLa cells transfected with hGPC4 (red curve) or with an irrelevant cDNA (cDNActrl; green curve). Cells were incubated with increasing concentrations of RB1-Fc. Data are presented as mean±SEM of n=3 biological replicates. Before pooling, data were normalized by the values of the negative control obtained with a an irrNb. Apparent K_D_ were calculated using the GraphPad Prism version 8 software. (**C**) qRT-PCR analyses of *hGPC4* mRNA levels in HeLA, MKN, SNU-449 cancer cells, in hiPSCs and hiPSCs with reduced *hGPC4* levels. Levels of all transcripts were normalized to those of the housekeeping gene GAPDH and subsequently to the ΔCt of the A549 cell lines that do not express *hGPC4*. Note the different *hGPC4* mRNA levels in cell lines. Data are presented as mean±SEM of n=2 biological replicates. One- way ANOVA with Dunnett’s multiple comparison test: \*\*\**p<0.*001. (**D**) Flow cytometry analysis of RB1-Fc cell binding on HeLa, MKN, SNU-449 cancer cells, on hiPSCs and hiPSCs with reduced hGPC4 levels. HGPC4 expression levels in different cell lines were evaluated by comparing the fluorescence staining given by RB1-Fc with that of an irrelevant Nb-Fc (irrNb-Fc). Note that despite low level of mRNA in SNU-449, RB1- Fc exhibits comparable binding to that observed in MKN cells, indicating similar surface target availability. (**E**) Flow cytometry analysis of RB1-Fc binding to hiPSCs and hiPSCs with reduced hGPC4 levels applied at 133nM concentrations. Data are presented as mean±SEM of n=3 biological replicates. Before pooling, data were normalized by the values of the negative control obtained with an irrelevant Nb-Fc. (**F**) Immunofluorescence analysis of hGPC4 in different cell lines using RB1-Fc. An irrNb- Fc was used as negative control. RB1-Fc detected hGPC4 in HeLa cells transfected with hGPC4 (HeLa + GPC4) as well as endogenous hGPC4 in cells such as MKN and hiPSCs producing relatively high hGPC4. No staining was detected in HeLa cells or hiPSCs GPC4sh due to their lack or low *hGPC4* transcript levels (see panel (C)). A commercially available mGPC4Ab was used as control of GPC4 expression on these cells.

The hGPC4-binding properties of the bivalent RB1-Fc was tested using different approaches. First, we determined the apparent affinity constant of RB1-Fc on hGPC4- transfected HeLa cells (Fig. 2B) by flow cytometry. Quantitative analysis from independent experiments showed a ∼14-fold increase in apparent binding affinity of RB1-Fc Nb over the monovalent Nb with a K_D_ shift from 84±73.16nM to 5.8±1.9 nM.

Next, we evaluated the ability of the RB1-Fc Nb to bind and detect various levels of endogenously expressed hGPC4. Using qRT-PCR analysis (Figure 2C), we selected a panel of cell lines based on hGPC4 mRNA expression levels: (i) high expression in MKN-45 cells (20,000 ± 657.1) and human induced pluripotent stem cells (hiPSCs; 16,108 ± 1,290), and (ii) moderate to low expression in hiPSCs GPC4sh (1,970 ± 110.1) and SNU-449 cells (1,395 ± 64.6) (Figure 2C). HeLa cells were used as negative control cell line. The binding capacity of RB1-Fc was assessed by flow cytometry at a Nb concentration of 44 nM. RB1-Fc bound to hGPC4 on all tested cell lines, including hiPSCs GPC4sh cells, which express markedly lower levels of GPC4 compared to hiPSC. This indicates that RB1-Fc is capable of detecting varying levels of hGPC4 expression (Figure 2C to 2E).

RB1-Fc binding to all cells was also tested by immunofluorescence. RB1-Fc successfully detected both exogenous (hGPC4-HeLa) and endogenous (MKN) GPC4 (Fig. 2F). Interestingly, although RB1-Fc binding was detectable on hiPSCs GPC4sh by flow cytometry – despite their low levels of membrane-bound GPC4 – no binding was observed on these cells by immunofluorescence.

### RB1-Fc recognizes a conformational epitope of the native hGPC4

The results described above demonstrate that RB1-Fc can interact with the cell-expressed hGPC4 protein. To study further the molecular interactions between RB1- Fc and hGPC4, we tested the RB1-Fc binding by western blot and immunoprecipitations.

Western blot analysis was performed on lysates from HEK cells transfected with human GPC4 (hGPC4) bearing a C-terminal HA tag. While the commercially available anti-HA antibody confirmed GPC4 expression, the RB1-Fc fusion protein did not detect any specific band. This indicates that RB1 does not recognize hGPC4 under reducing and denaturing conditions (Figure S4A).

Next, we performed immunoprecipitation experiments using lysates from HEK cells transfected with hGPC4 constructs containing an HA tag either at the C-terminus as above or at the N-terminus. These experiments were performed under both non-denaturing and denaturing conditions. The use of both N- and C-terminally tagged constructs allowed us to control for potential interference of the HA tag with RB1-Fc binding to hGPC4. RB1-Fc successfully immunoprecipitated hGPC4 when cells were lysed by non denaturing buffers (Figure 3A and Figure S4B). In contrast, the use of denaturing lysis buffers containing the harsh ionic detergent SDS – known to disrupt protein structure and charge – drastically impaired the ability of RB1-Fc to immunoprecipitate cell-expressed hGPC4 (Figure 3B). These findings indicate that RB1-Fc interacts with hGPC4 only in its native conformation. Notably, RB1-Fc was able to immunoprecipitated both the glycanated forms [9–11] and the core protein of hGPC4 (Figure 3A), suggesting that binding of RB1-FC to hGPC4 does not depend on the presence of the heparan sulphate side chains.

**Figure 3.**
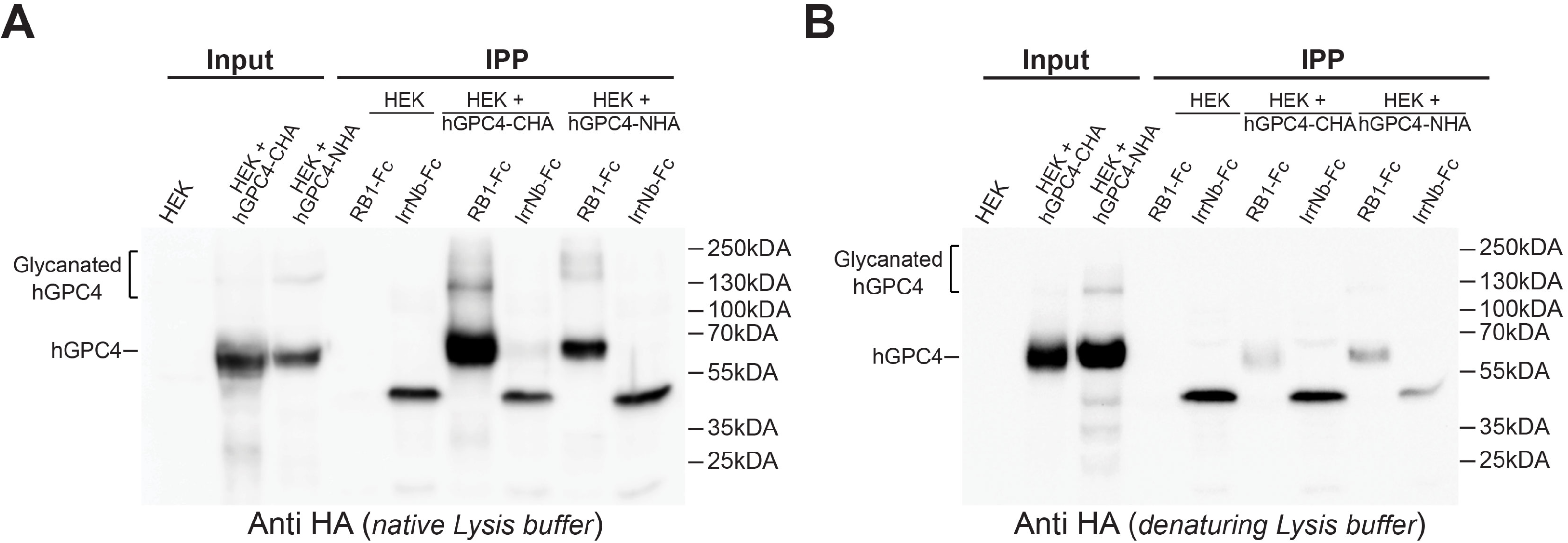
RB1-Fc targets a conformational epitope of hGPC4. (**A-B**) Immunoprecipitation assay with cell extracts from non-transfected HEK cells (HEK) or transfected with a hGPC4 with an HA tag at the carboxyterminal (hGPC4-CHA) or a hGPC4 with an HA tag at the amminoterminal (hGPC4-NHA). Protein extracts were prepared using non denaturing buffer 1 (**A**) or denaturing (**B**) buffers and analysed by SDS-PAGE under reducing conditions. Immunoprecipitations were performed using RB1-Fc, and irrNb-Fc. Immunoprecipitated proteins were detected by western blot using anti HA antibodies. Note that the hGPC4 was immunoprecipitated from native cell lysates incubated with RB1-Fc and not with the irrNb-Fc control (A). The immunoprecipitation of hGPC4 from a denatured cell lysate was found to be inefficient (B). The asterisks in (A) indicate the glycanated hGPC4 forms. Note that the band of approximately 50 kDa detected in the lanes corresponding to irrNb-Fc represents the irrNb-Fc nanobody, as it is recognized by the anti-HA antibody targeting its HA tag epitope.

### RB1-Fc blocks hGPC4 biological functions

As discussed above, impairing hGPC4 functions in hiPSCs and in cancer cell types - such as pancreatic cancer stem cells - can potentially provide a clinically relevant strategy to both enhance the therapeutic potential of hiPSCs and target specific cancers. In this context, the development of blocking antibodies capable of inhibiting hGPC4 activity would represent a valuable tool for both regenerative medicine and oncology applications. In the perspective of using RB1-Fc as hGPC4 blocking reagent, we first established whether RB1-Fc would elicit toxic effects on cells. Our previous loss-of-function studies in hiPSCs have shown that downregulation of hGPC4 in these cell types does not affect their survival [25, 28]. Therefore, RB1-Fc toxic effects were determined by exposing hiPSCs to increasing concentration of RB1-Fc, using an irrelevant Nb-Fc as negative control. Quantification analysis of surviving cells, done using a metabolic activity–based cell viability assay, revealed that RB1-Fc did not elicit significant cytotoxic effects even at concentrations up to 1000 nM (Figure 4A).

**Figure 4.**
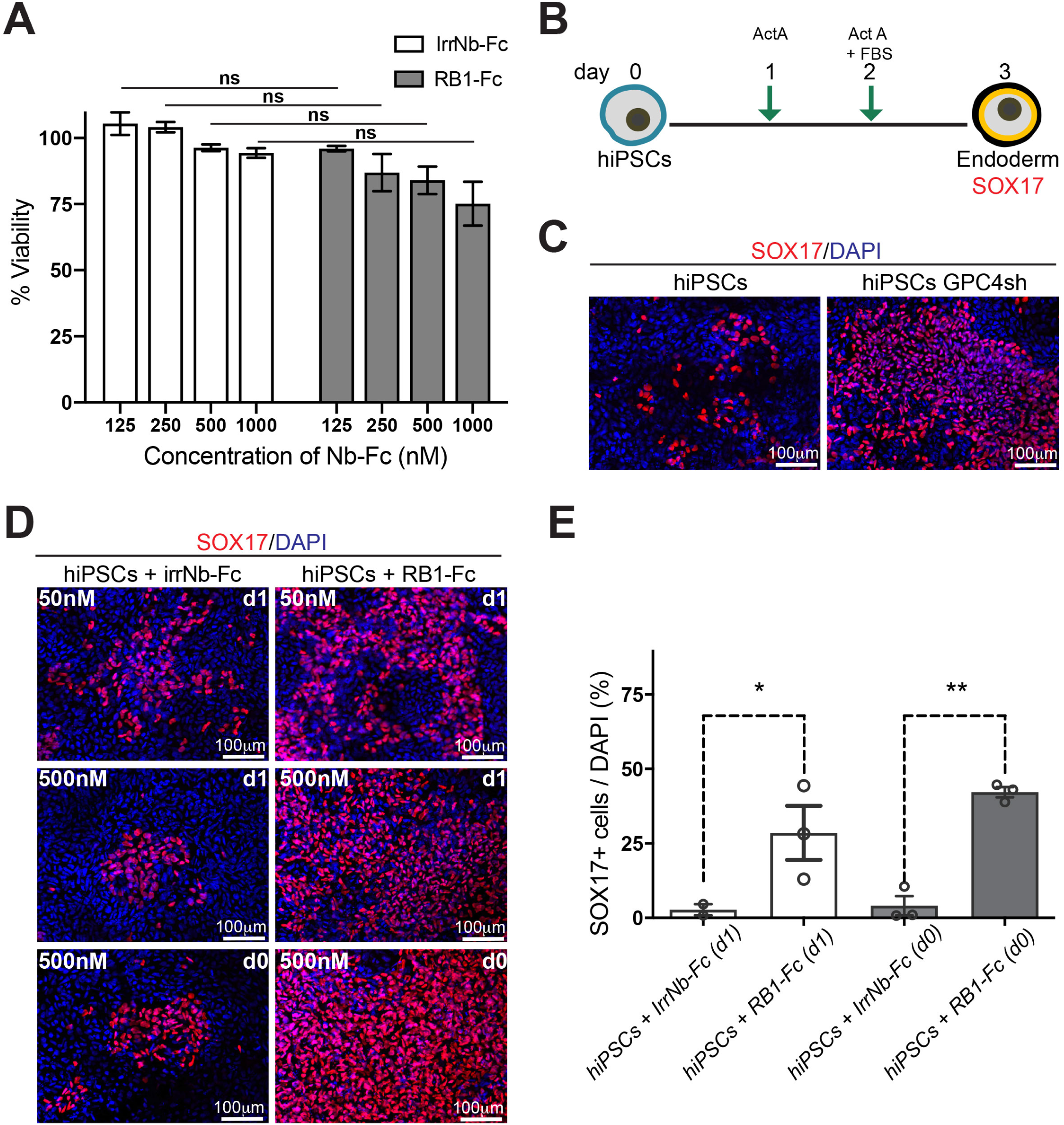
RB1-Fc elicits hGPC4-blocking activity. (**A**) Viability of hiPSCs in the presence of increasing concentrations of RB1-Fc and irrNb-Fc. Percentage of viable cells was measured in a metabolic activity-based cell viability assay. Data are presented as mean±SEM of n=4 biological replicates. One-Way ANOVA with Tukey’s multiple comparison tests: \*\*\**p<0.*001. Note that not significant RB1-Fc toxic effects were observed at concentrations up to 1000nM. (**B**) Schematic representation of the endoderm differentiation protocol of hiPSCs applied to test the RB1-Fc blocking activity on hGPC4. hiPSCs are plated at day 0 and exposed to ACTIVIN A (ActA) and ACTIVIN A + 0.2% FBS (ActA+FBS) at day 1 and day 2 of differentiation, respectively. Endoderm differentiation is analyzed at day 3 by following the number of SOX17 positive cells. (**C**) Representative images of SOX17 positive cells in hiPSCs and hiPSCs with reduced hGPC4 levels (hiPSCs GPC4sh) after 3 days of differentiation showing the increased distribution of endodermal cells in the hiPSCs GPC4sh line. (**D**) Representative images of SOX17 positive cells in hiPSCs exposed to irrNb-Fc control and to RB1-Fc at 50 and 500nM. hiPSCs were treated with the Nbs either from day1 of differentiation or starting from day 0. Note that the increased distribution of SOX17 positive cells in hiPSCs treated with RB1-Fc is comparable to that observed in the hiPSCs GPC4sh line (C). (**E**) Quantitative analysis of SOX17 positive cells in hiPSCs treated with a control (irrNb-Fc) and with RB1-Fc at 500 nM. SOX17-positive cells were defined as those whose mean fluorescence intensity was above the 95^th^ percentile of the mean SOX17 intensity in negative cells at the same time point. At least 10 randomly selected frames of view were taken per condition, covering >2.23 mm² in total. Data are presented as mean±SEM of n=3 biological replicates. One-Way ANOVA with Tukey’s multiple comparison tests: \*\**p<0.*01. Note a ∼10 % increase of SOX17 positive cells in differentiating hiPSCs incubated with RB1-Fc in comparison to hiPSCs incubated with an irrelevant Nb. Also, hiPSCs exposed to RB1-Fc at day 0 have a higher trend for generating SOX17 positive cells in comparison to cells treated with RB1-Fc starting from day1.

Next, we examined whether RB1-Fc could function as hGPC4 blocking reagent. We have recently established that hiPSCs in which hGPC4 proteins levels are reduced by means of shRNA targeting (GPC4sh) undergo a more efficient endoderm lineage entry than controls in response to endoderm triggering factors [28]. This phenotype is apparent upon three-day treatment of hiPSCs with ACTIVIN-A (Figure 4B) as evidenced by a ∼17-fold increase in the number of SOX-17 positive cells detected by immunocytochemistry (Figure 4C). We, therefore, tested whether treatment of hiPSCs with the RB1-Fc impairs hGPC4 activity, leading to a more efficient endoderm lineage entry than hiPSCs GPC4sh. The ability of RB1-Fc to influence the endoderm differentiation of hiPSCs was tested by performing differentiation experiments in the presence of different concentration of RB1-Fc, irrelevant Nb-Fc being used as negative control. Nb-Fc fusions were applied at the concentrations of 50 and 500nM, as they do not cause toxic effects on cells (Figure 4A). Two different settings were compared. In the first setting, RB1-Fc was applied at the onset of endoderm differentiation (d1; Figure 4B) and maintained throughout the differentiation process (d1-d3; Figure 4B). In the second setting, hiPSCs were pre-treated with RB1-Fc for 24 hours prior to the induction of differentiation (day 0), and RB1-Fc treatment continued during the entire differentiation period (days 0–3; Figure 4B). The rationale for this second method was based on the assumption that blocking hGPC4 functions in undifferentiated hiPSCs would confer biological properties more similar to those of hiPSCs GPC4sh [28], thus enabling a more efficient response to differentiation signals. In both protocols, treatment of hiPSC with RB1-Fc drastically increased their ability to enter the endodermal lineage as revealed by the abundance of SOX-17 positive cells (Figure 4D). A quantitative analysis demonstrated that treatment with 500 nM RB1-Fc resulted in at least ∼20-fold increase of SOX17 positive cells during differentiation of hiPSCs, compared to cells treated with the same concentration of an irrelevant Nb (Figure 4E). This increase in SOX17-positive cells was comparable with that observed in hiPSCs GPC4sh (Figure 4C; [28]), Interestingly, hiPSCs exposed to RB1-Fc for 24 hours prior to the onset of differentiation (d0) exibited a greater propensity to generate SOX17- positive cells compared to those treated at the start of differentiation (d1) (Figure 4D and 4E), suggesting that early RB1-Fc treatment potentiates its antagonist efftect on hGPC4 functions. Altogether, these results define the Nb RB1-Fc as the first hGPC4- blocking antibody offering a novel strategy to inhibit hGPC4 activity without the need for genetic manipulations. They also suggest that RB1-Fc can serve as a valuable tool to study hGPC4 functions in a temporal and spatial controlled manner.

## DISCUSSION

The fine-tuning of environmental signals provides the basis for proper development and physiological processes[48]. Glypicans have emerged as key modulators of cell communication across a wide range of biological contexts, and mutations in glypican genes have been implicated in several human pathologies [9–11, 15]. This has sparked significant interest within the biomedical research community, which is actively exploring their potential as diagnostic and prognostic markers, as well as therapeutic targets. Nanobodies – distinguished from conventional antibodies by their small size, high affinity and specificity, and ability to access cryptic epitopes – have recently gained prominence as powerful tools for studying and modulating target antigens in living cells [42].

In the present work, we focused on the generation of nanobodies targeting hGPC-4, a member of glypican family that has emerged as both biomarker and a potential therapeutic target in several human diseases [31–36]. HGPC-4 has also been identified as a key target for regulating hPSC biology and enhancing their therapeutic potential [25–28]. Here, we report for the first time the generation of two hGPC4-specific single domain antibodies, RB1 and RB3. While RB3 binds exclusively to the recombinant hGPC-4 protein, RB1 recognizes both recombinant and cell surface-associated hGPC-4. As expected from its bivalent structure, the RB1-Fc fusion construct significantly increases the apparent binding affinity of RB1 to membrane-bound hGPC4 – by approximately 14-fold – enabling the specific detection of cells expressing varying levels of endogenous hGPC4. Furthermore, we demonstrated that RB1-Fc binds hGPC4 only in its native conformation, suggesting that RB1-Fc recognizes a conformational epitope.

Although the exact RB1-Fc binding site on hGPC4 remains to be determined, our biochemical analyses suggest that it is localized on the core protein of hGPC4 rather than on the heparan sulfate chains, as shown by its ability to immunoprecipitate both the glycanated forms and core protein of hGPC4. This, along with the recognition of a conformational epitope, highlights RB1-Fc as a high-affinity, specific binder of native, cell surface-associated hGPC4. Future structural studies will be essential to precisely define the epitope recognized by RB1-Fc.

The specificity of the RB1-Fc Nb for hGPC4 is further supported by functional studies in hiPSCs, which demonstrated its blocking activity. Differentiation analyses revealed that RB1-Fc–treated hiPSCs exhibited a significantly enhanced capacity to differentiate into endoderm, closely resembling the phenotype observed in hiPSCs with downregulate hGPC4. These results suggest that RB1-Fc binding functionally inhibits hGPC4, potentially acting as an orthosteric competitor or an allosteric negative modulator. These findings support the notion that pharmacological targeting of hGPC4 with RB1-Fc represents a simple and effective alternative to genetic manipulation for promoting hiPSC differentiation. RB1-Fc may therefore help overcoming current limitations in stem cell research and promote the broader application of hPSCs in disease modeling, drug screening, and regenerative medicine.

The growing body of evidence linking alteration in hGPC-4 protein levels to metabolic and CNS disorders – such as high body fat content, insulin resistance and Alzheimer’s diseases – underscores the need to accurately quantify cell-surface antigen density in clinical samples. Our findings demonstrate that RB1-Fc is a promising tool for detecting changes in hGPC-4 protein levels. While commercial ELISA kits are available for the detection of hGPC-4 in biological fluids, concerns have been raised regarding their specificity and suitability for accurately measuring hGPC-4 [49]. In contrast, our experiments highlight both sensitivity and specificity of RB1-Fc for hGPC-4 detection. Furthermore, due to the modular nature of Nbs, further development of higher- valencies – such as hexavalent format consisting of three tandem of RB1 units dimerised via Fc fusion – could enhance binding strength and detection capabilities.

In addition to metabolic and neurological disorders, hGPC4-targeting nanobodies like RB1-Fc may have broad clinical applications. One promising area is cancer diagnosis and therapy, as increasing evidence suggests a correlation between elevated circulating hGPC4 levels and poor prognosis in glioblastoma, colorectal cancer, and metastatic breast cancer[31, 32]. To our knowledge, there are no well-established small molecules or chemical inhibitors currently available that specifically target hGPC4. Molecules such as surfen or heparin mimetics are designed to disrupt heparan sulfate (HS)–protein interactions and modulate HS-dependent signaling [50]. However, their lack of specificity often results in widespread effects on multiple HS-mediated pathways, potentially leading to unwanted impacts on diverse cellular functions. In some systems, a soluble hGPC4 ectodomain is used as a decoy to compete with endogenous GPC4 for ligand binding [51]. This does not block hGPC4 per se but modulates its pathways. Most previous studies on hGPC4 inhibition in cells and animal models relied on siRNA or shRNA-mediated knockdown [25, 52]. In this context, RB1- Fc represents the first antibody reported to directly block hGPC4 activity. With the FDA approval of caplacizumab in 2019 as the first therapeutic nanobody, numerous nanobody-based agents are now advancing through clinical development[53]. It will be of great interest to explore whether RB1-Fc could become a clinical candidate for targeting hGPC4-dependent cancers. Notably, loss of hGPC4 activity in pancreatic cancer cells has been shown to sensitize these cells to chemotherapeutic agents and reduce stem cell-like properties [30]. In this context, hGPC4 modulators such as RB1- Fc may enhance the effectiveness of existing chemotherapy regimens.

Beyond its diagnostic and therapeutic potential, RB1-Fc is expected to serve as a valuable tool for basic research into hGPC4 function in both embryonic and adult cells. For example, proteomic approaches leveraging RB1-Fc could facilitate the identification of cytokines, morphogens, and other proteins that interact with hGPC4 across various cellular contexts. Such studies may uncover novel hGPC4-mediated mechanisms of action and provide deeper insight into its roles in development and disease.

## AUTHOR CONTRIBUTIONS

**Remi Bonjean**: Writing – review & editing, Conceptualization, Methodology, Investigation, Data curation, Formal analysis. **Brigitte Kerfelec**: Writing – review & editing, Data curation. **Patrick Chames**: Writing – review & editing, Conceptualization, Methodology, Data curation. **Rosanna Dono**: Writing – original draft, Writing – review & editing, Conceptualization, Methodology, Investigation, Data curation, Formal analysis, Project administration, Funding acquisition

## ACKNOWLEDGEMENTS

We thank: all members of our previous and current labs for helpful discussions and comments; Daniel Baty for his help and advice at the start of this project. T. Legier, D. Rattier for their assistance in conducting the initial studies on hiPSC differentiation and cell viability. Microscopy was performed at the imaging platform of the IBDM, supported by the ANR through the "Investments for the Future" program (France-BioImaging, ANR-10-INSB-04-01). The authors thank members of the IBDM core microscopy facility for their technical support.

## DATA AVAILABILITY

Data will be made available on request

The authors declare no competing or financial interests.

---

**Figure S1.**
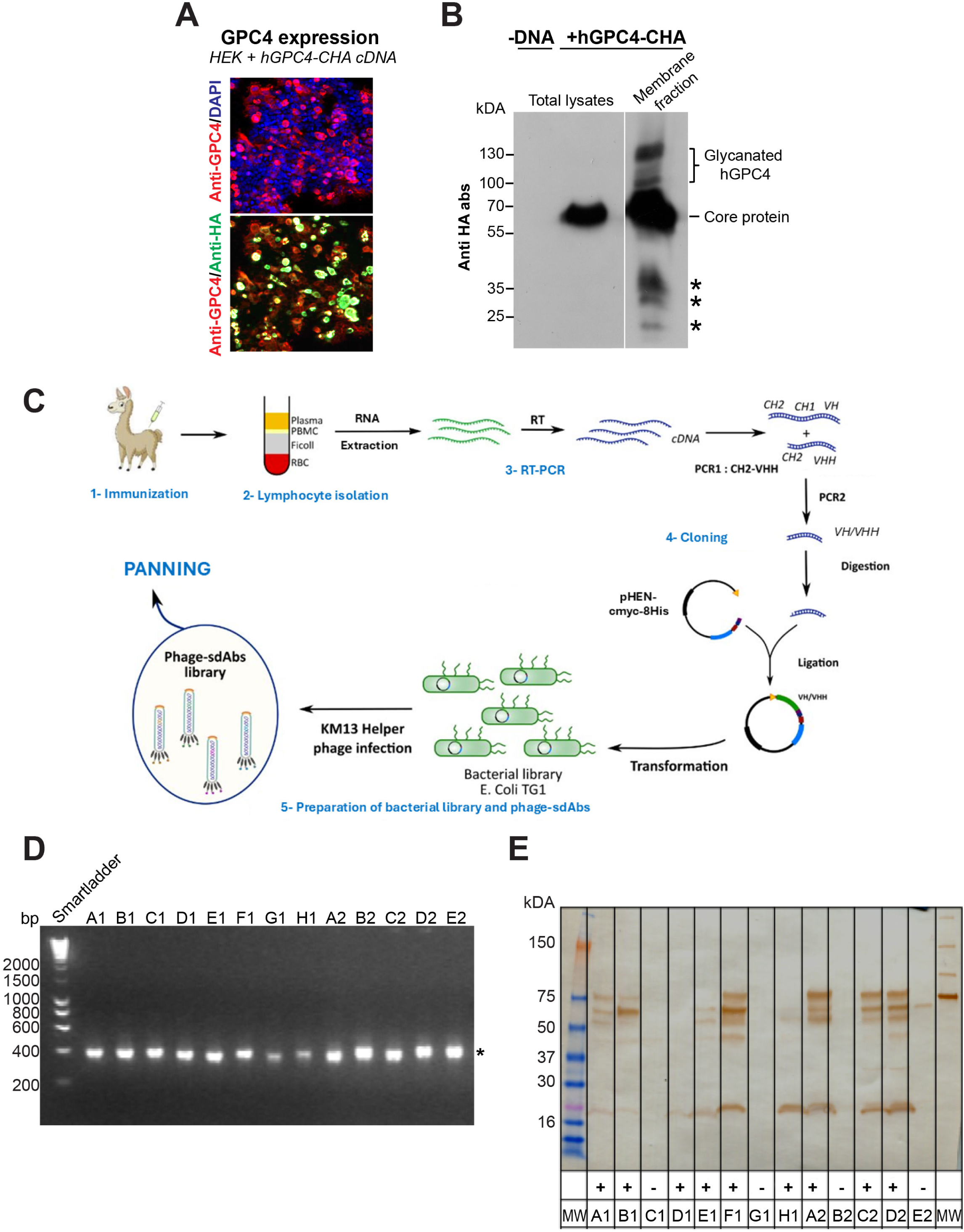
Preparation of the protein extract for llama immunization and phage display libraries. (**A**) Immunofluorescence analysis of HEK cells transfected with the hGPC4 protein fused to a carboxyterminal HA-tag (hGPC4-CHA). (**B**) Western blot analysis showing exogenous hGPC4 in the total cell lysate and in the membrane fraction used for llama immunization. Note the core protein and all glycanated hGPC4 forms. The N-terminal and C-terminal hGPC4 fragments, generated during or after hGPC4 synthesis and processing are also visible (asterisks) [10]. (**C**) Schematic procedure for the production of phage display Nb libraries. Blood samples were collected at day 28 and day 42 for lymphocyte preparation. (**D**) PCR analysis of phage display library transformants showing that they harbor a vector with an insert of the VHH size. (**E**) Supernatant of individual phage clones analyzed by Western blot. Lane MW: molecular weight marker. Lanes A1 to E2 Nb proteins. The + and – symbols indicated colonies expressing or not the Nb proteins.

**Figure S2.**
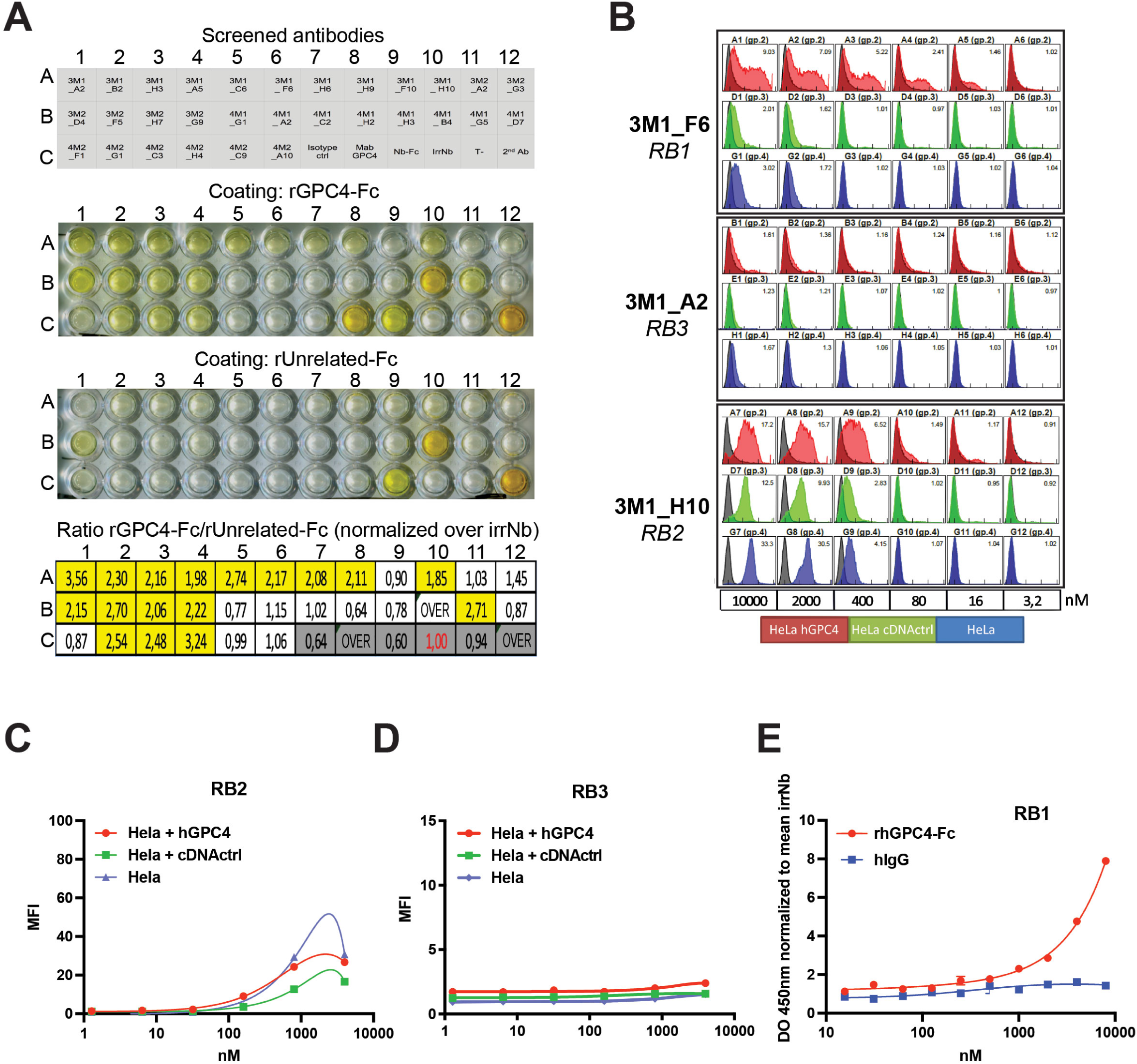
Identification of GPC4-targeting Nbs. (**A**) Representative results of the GPC4-targeting Nb-screening done by ELISA. Supernatant of individual phage clones (top scheme) was screened by ELISA on plates coated with recombinant hGPC4-Fc (Coating: rGPC4-Fc) or with unrelated Fc tagged recombinant protein as negative control (Coating: rUnrelated-Fc). Values were normalized over those obtained with an irrNb (well C10) and reported as ratio between rGPC4-Fc/rUnrelated-Fc. Positive control: mAb GPC4. Negative controls: Isotype ctrl, irrNb-Fc (Nb-Fc), Irr-Nb, T-, 2^nd^ Ab. (B) Representative results of the hGPC4-targeting Nb-screening done by Flow cytometry. Nbs were tested for binding properties to HeLA cells expressing hGPC4 (HeLA hGPC4, red curves), HeLA cells transfected with a non-specific plasmid (HeLA cDNA CTRL, green curves) and not transfected cells (HeLA, blue curves). Nbs were added at increasing concentrations (nM). Note that the Nb 3M1_F6 thereafter named as RB1 bind to cells expressing hGPC4. In contrast, 3M1_A2, thereafter named as RB3, does not bind to cells expressing hGPC4. The Nb 3M1_H10, thereafter named as RB2, turned out as not specific hGPC4-binder as it recognizes negative controls. (C) FACS analysis of cell binding on HeLa cells non-transfected (blue curve) or transfected with hGPC4 (red curve) or with an irrelevant cDNA (cDNActrl; green curve). Cells were incubated with increasing concentrations of RB2. n=1 biological replicates. (D) FACS analysis of cell binding on HeLa cells non-transfected (blue curve) or transfected with hGPC4 (red curve) or with an irrelevant cDNA (cDNActrl; green curve). Cells were incubated with increasing concentrations of RB3. n=1 biological replicates. (E) Binding of RB1 to recombinant hGPC4-Fc protein (red curve) and to hIgGs (blue curve) examined by ELISA. RB1 was added at increasing concentrations. Data are presented as mean±SEM of n=2. Before pooling, data were normalized by the values of the negative control obtained with a an irrNb. Note that RB1 binds to recombinant hGPC4-Fc protein with low affinity (Kd apparent >4000nM).

**Figure S3.**
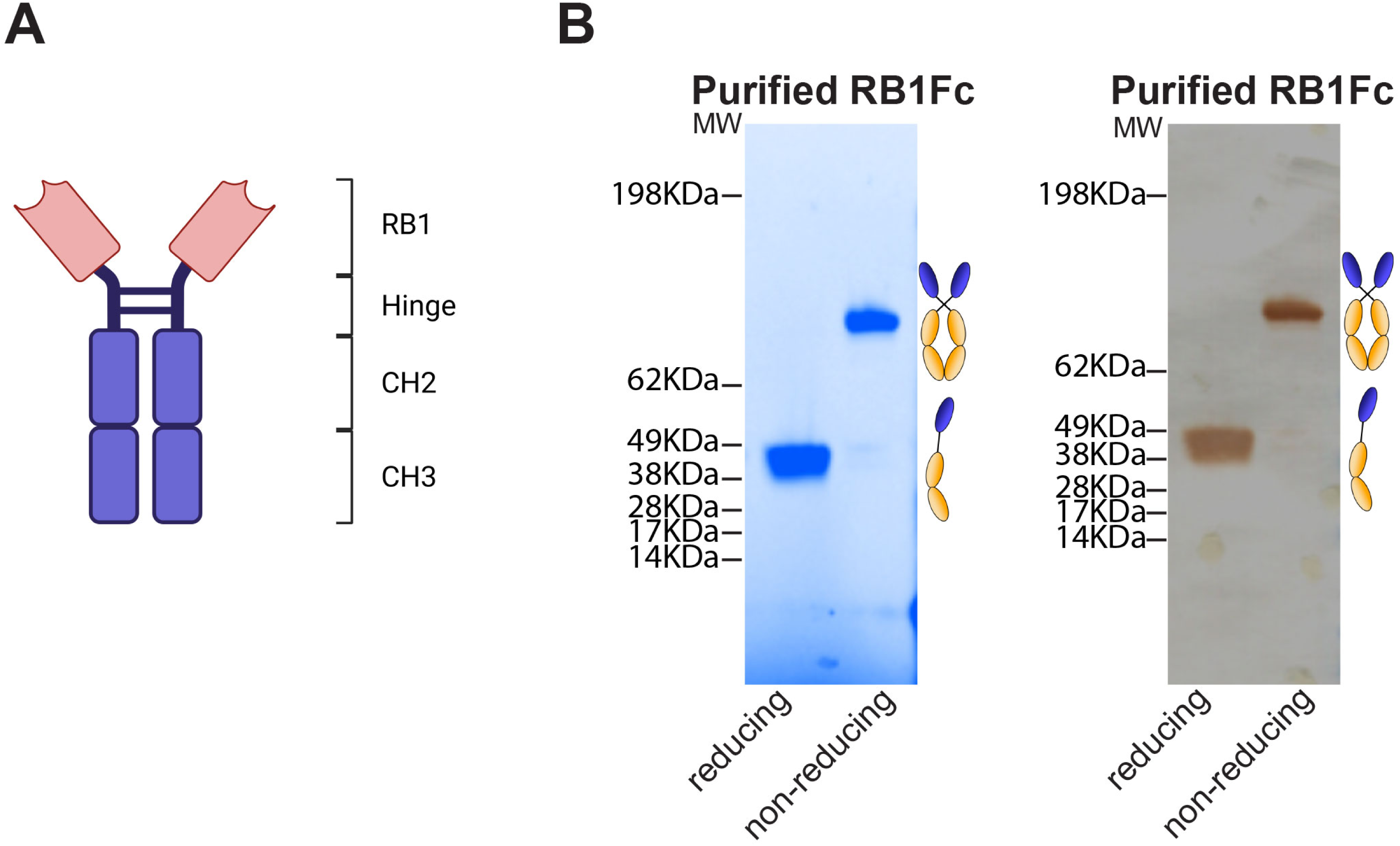
Generation of RB1-Fc. (**A**) Scheme representing the RB1-Fc fusion protein. (**B**) Analysis of the purified RB1-Fc Nb by SDS-PAGE (left) and western blot (right) shows the reducing and non-reducing RB1-Fc forms (right schemes).

**Figure S4.**
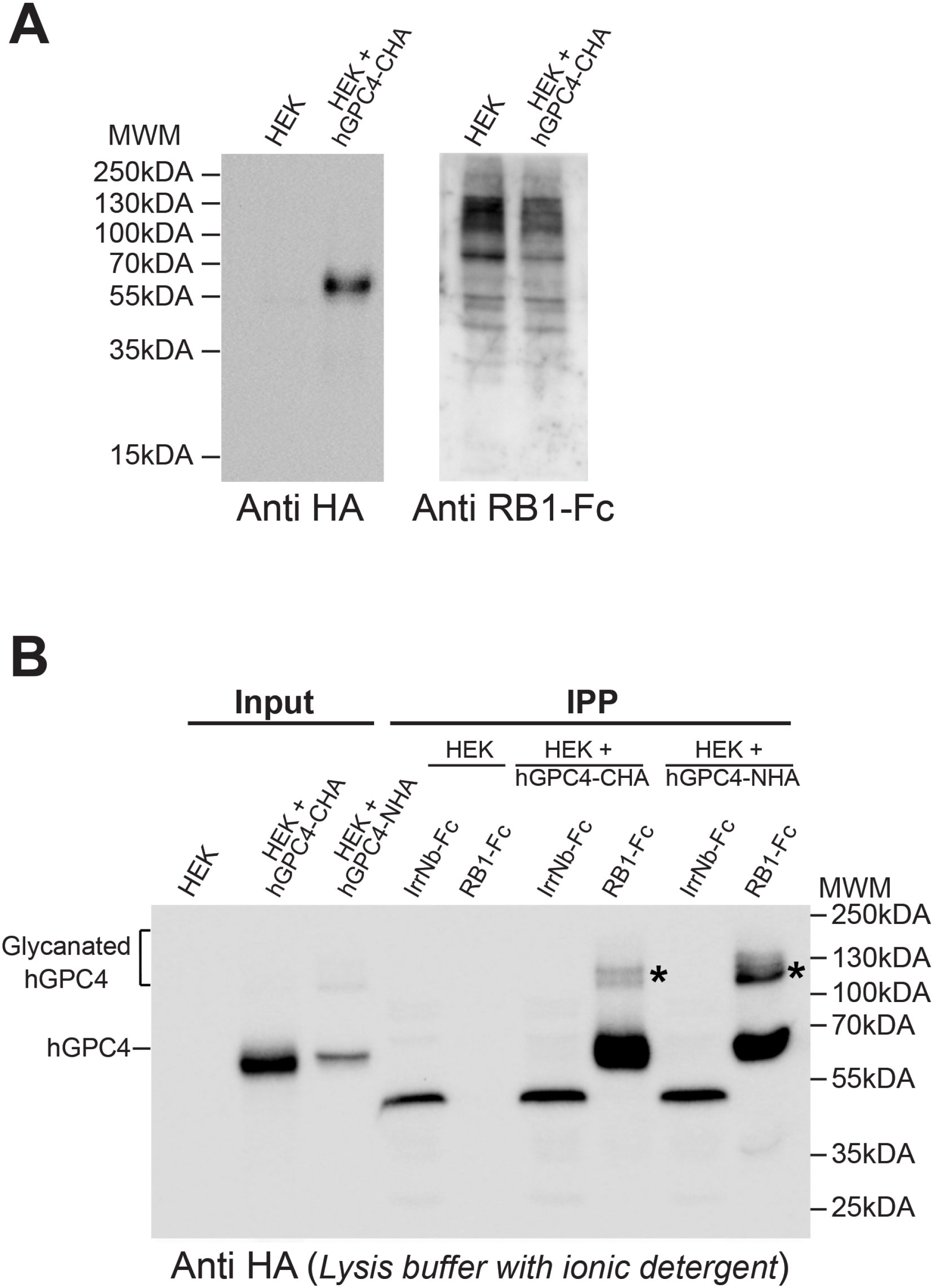
RB1-Fc targets a conformational epitope of hGPC4. (**A**) Western blot analysis of cell extracts from non-transfected HEK cells (HEK), transfected with a hGPC4 with an HA tag at the carboxyterminal (HEK + hGPC4-CHA). Proteins were detected by western blot using anti HA antibodies (left) and RB1-Fc (right) followed by HRP-conjugated anti-IgG. Note that the anti HA antibodies succesfully detected the transfected hGPC4-CHA protein. In contrast, RB1-Fc did not reveal any specific band when compared to lysate from non-transfected HEK cells. The observed bands in the RB1-Fc western blot are considered non-specific signal due to FB1-Fc. (**B**) Immunoprecipitation assay with cell extracts from non-transfected HEK cells (HEK), transfected with a hGPC4 with an HA tag at the carboxyterminus (hGPC4-CHA) or a hGPC4 with an HA tag at the amminoterminus(hGPC4-NHA). Protein extracts were prepared using a lysis buffer with the ionic detergent, sodium deoxycholate (non denaturing buffer 2). Immunoprecipitations were performed using RB1-FC, an irrNb- Fc. Immunoprecipitated proteins were detected by western blot using anti HA antibodies. Note that the hGPC4 was immunoprecipitated from native cell lysates incubated with RB1-Fc and not with the irrNb-Fc control. The asterisks in (B) indicate the Glycanated hGPC4 forms. Note that the band of approximately 50 kDa detected in the lanes corresponding to irrNb-Fc represents the irrNb-Fc nanobody, as it is recognized by the anti-HA antibody targeting its HA tag epitope.

**Table S1.** List of all primary and secondary antibodies used. ICC=Immunocytochemistry, ELISA=E, WB= Western Blot, FL=Flow Cytometry.

| ANTIGEN | SPECIES | Conjugated | COMPANY | REF. | DILUTION |
| --- | --- | --- | --- | --- | --- |
| 6His tag | Mouse | HRP | Milteniy Biotec | 130-092-785 | 1:5000 (E) |
| 6His tag | Mouse |  | Merck Millipore | 7096-3 | 1:1000 (FC) |
| Fc tag | Goat | HRP | Sigma-Aldrich | A0170 | 1:5000 (E; WB) |
| Fc tag | Goat | Phycoérythrine | Beckman | IM0550 | 1:250 (FC) |
| GPC4 | Mouse |  | Genetex | AT4A3 | 1:200 (E; ICC) |
| Flag tag | Mouse |  | Sigma-Aldrich | F3165 | 1:1000 (FC) |
| HA tag | Rat |  | Roche | 11867431<br>001 | 1:500<br>(ICC)<br>1:1000<br>(WB) |
| SOX17 | Goat |  | R&D | 967330 | 1:100<br>(ICC) |
| Anti-Rat | Donkey | HRP | Jackson | 712035150 | 1:4000<br>(WB) |
| Anti-hFc | Goat | HRP | ThermoFischer | A18817 | 1:2000 (E;<br>WB) |
| Anti-Mouse | Goat | HRP | SantaCruz | SC-2005 | 1:5000 (E) |
| Anti-Mouse | Goat | Alexa647 | Invitrogen | A21235 | 1:500 (FC) |
| Anti-human | Goat | HRP | Jackson | 109-035-088 | 1:5000 (E) |
| Anti-goat | Donkey | Alexa647 | Invitrogen | A-21447 | 1:500<br>(ICC) |
| Anti-rat | Donkey | Alexa647 | Invitrogen | A48272 | 1:500<br>(ICC) |

## REFERENCES

[1] M. Freeman, J.B. Gurdon, Regulatory principles of developmental signaling, Annu Rev Cell Dev Biol 18 (2002) 515–39.

[2] P. Muller, A.F. Schier, Extracellular movement of signaling molecules, Dev Cell 21(1) (2011) 145–58.

[3] B.E. Housden, N. Perrimon, Spatial and temporal organization of signaling pathways, Trends Biochem. Sci 39(10) (2014) 457–64.

[4] M. Majidinia, A. Sadeghpour, B. Yousefi, The roles of signaling pathways in bone repair and regeneration, J Cell Physiol 233(4) (2018) 2937–2948.

[5] N. Perrimon, C. Pitsouli, B.Z. Shilo, Signaling mechanisms controlling cell fate and embryonic patterning, Cold Spring Harb Perspect Biol 4(8) (2012) a005975.

[6] Z. Steinhart, S. Angers, Wnt signaling in development and tissue homeostasis, Development 145(11) (2018).

[7] S.G. Wilcockson, C. Sutcliffe, H.L. Ashe, Control of signaling molecule range during developmental patterning, Cell Mol Life Sci 74(11) (2017) 1937–1956.

[8] U. Hacker, K. Nybakken, N. Perrimon, Heparan sulphate proteoglycans: the sweet side of development, Nat Rev Mol Cell Biol 6(7) (2005) 530–41.

[9] F.E. Poulain, H.J. Yost, Heparan sulfate proteoglycans: a sugar code for vertebrate development?, Development 142(20) (2015) 3456–67.

[10] A. Fico, F. Maina, R. Dono, Fine-tuning of cell signaling by glypicans, Cellular and Molecular Life Sciences 68(6) (2011) 923–929.

[11] A.a.D. Fico, R., Signalling mechanisms underlying congenital malformation: the gatekeepers, glypicans., Congenital Malformations., Alastair Sutcliffe, University College London, University of London, United Kingdom 2011.

[12] I. Matsuo, C. Kimura-Yoshida, Extracellular distribution of diffusible growth factors controlled by heparan sulfate proteoglycans during mammalian embryogenesis, Philos Trans R Soc Lond B Biol Sci 369(1657) (2014).

[13] D. Yan, X. Lin, Shaping morphogen gradients by proteoglycans, Cold Spring Harb Perspect Biol 1(3) (2009) a002493.

[14] S. Kakugawa, P.F. Langton, M. Zebisch, S. Howell, T.H. Chang, Y. Liu, T. Feizi, G. Bineva, N. O’Reilly, A.P. Snijders, E.Y. Jones, J.P. Vincent, Notum deacylates Wnt proteins to suppress signalling activity, Nature 519(7542) (2015) 187–192.

15. [15] J. Filmus, Glypicans, 35 years later, Proteoglycan Research 1 (2023) e5.

[16] A. Galli, A. Roure, R. Zeller, R. Dono, Glypican 4 modulates FGF signalling and regulates dorsoventral forebrain patterning in Xenopus embryos, Development 130(20) (2003) 4919–29.

[17] B.D. Jakubovic, S. Jothy, Glypican-3: from the mutations of Simpson-Golabi-Behmel genetic syndrome to a tumor marker for hepatocellular carcinoma, Exp Mol Pathol 82(2) (2007) 184–9.

[18] N. Li, W. Gao, Y.F. Zhang, M. Ho, Glypicans as Cancer Therapeutic Targets, Trends Cancer 4(11) (2018) 741–754.

[19] J. Topczewski, D.S. Sepich, D.C. Myers, C. Walker, A. Amores, Z. Lele, M. Hammerschmidt, J. Postlethwait, L. Solnica-Krezel, The zebrafish glypican knypek controls cell polarity during gastrulation movements of convergent extension, Dev Cell 1(2) (2001) 251–264.

[20] D. Li, S. Lin, J. Hong, M. Ho, Immunotherapy for hepatobiliary cancers: Emerging targets and translational advances, Advances in Cancer Research, Elsevier 2022, pp. 415–449.

[21] T. Nishida, H. Kataoka, Glypican 3-Targeted Therapy in Hepatocellular Carcinoma, Cancers (Basel) 11(9) (2019).

[22] S. Heitzeneder, K.R. Bosse, Z. Zhu, D. Zhelev, R.G. Majzner, M.T. Radosevich, S. Dhingra, E. Sotillo, S. Buongervino, G. Pascual-Pasto, E. Garrigan, P. Xu, J. Huang, B. Salzer, A. Delaidelli, S. Raman, H. Cui, B. Martinez, S.J. Bornheimer, B. Sahaf, A. Alag, I.S. Fetahu, M. Hasselblatt, K.R. Parker, H. Anbunathan, J. Hwang, M. Huang, K. Sakamoto, N.J. Lacayo, D.D. Klysz, J. Theruvath, J.G. Vilches-Moure, A.T. Satpathy, H.Y. Chang, M. Lehner, S. Taschner-Mandl, J.P. Julien, P.H. Sorensen, D.S. Dimitrov, J.M. Maris, C.L. Mackall, GPC2- CAR T cells tuned for low antigen density mediate potent activity against neuroblastoma without toxicity, Cancer Cell 40(1) (2022) 53–69 e9.

[23] N. Li, H. Fu, S.M. Hewitt, D.S. Dimitrov, M. Ho, Therapeutically targeting glypican-2 via single-domain antibody-based chimeric antigen receptors and immunotoxins in neuroblastoma, Proc Natl Acad Sci U S A 114(32) (2017) E6623–E6631.

[24] R. Dono, Glypican 4 down-regulation in pluripotent stem cells as a potential strategy to improve differentiation and to impair tumorigenicity of cell transplants, Neural Regen Res 10(10) (2015) 1576–7.

[25] S. Corti, R. Bonjean, T. Legier, D. Rattier, C. Melon, P. Salin, E.A. Toso, M. Kyba, L. Kerkerian-Le Goff, F. Maina, R. Dono, Enhanced differentiation of human induced pluripotent stem cells toward the midbrain dopaminergic neuron lineage through GLYPICAN-4 downregulation, Stem Cells Transl Med 10(5) (2021) 725–742.

[26] A. Fico, A. De Chevigny, J. Egea, M.R. Bosl, H. Cremer, F. Maina, R. Dono, Modulating Glypican4 suppresses tumorigenicity of embryonic stem cells while preserving self-renewal and pluripotency, Stem Cells 30(9) (2012) 1863–74.

[27] A. Fico, A. de Chevigny, C. Melon, M. Bohic, L. Kerkerian-Le Goff, F. Maina, R. Dono, H. Cremer, Reducing Glypican-4 in ES Cells Improves Recovery in a Rat Model of Parkinson’s Disease by Increasing the Production of Dopaminergic Neurons and Decreasing Teratoma Formation, Journal of Neuroscience 34(24) (2014) 8318–8323.

[28] T. Legier, D. Rattier, J. Llewellyn, T. Vannier, B. Sorre, F. Maina, R. Dono, Epithelial disruption drives mesendoderm differentiation in human pluripotent stem cells by enabling TGF-beta protein sensing, Nat Commun 14(1) (2023) 349.

[29] Y. Fang, Z.Y. Shen, Y.Z. Zhan, X.C. Feng, K.L. Chen, Y.S. Li, H.J. Deng, S.M. Pan, D.H. Wu, Y. Ding, CD36 inhibits beta-catenin/c-myc-mediated glycolysis through ubiquitination of GPC4 to repress colorectal tumorigenesis, Nat Commun 10(1) (2019) 3981.

[30] J. Cao, J. Ma, L. Sun, J. Li, T. Qin, C. Zhou, L. Cheng, K. Chen, W. Qian, W. Duan, F. Wang, E. Wu, Z. Wang, Q. Ma, L. Han, Targeting glypican-4 overcomes 5-FU resistance and attenuates stem cell-like properties via suppression of Wnt/beta-catenin pathway in pancreatic cancer cells, J Cell Biochem 119(11) (2018) 9498–9512.

[31] A. Muendlein, C. Heinzle, A. Leiherer, E.M. Brandtner, K. Geiger, S. Gaenger, P. Fraunberger, A. Mader, C.H. Saely, H. Drexel, Circulating glypican-4 is a new predictor of all-cause mortality in patients with heart failure, Clin Biochem 121–122 (2023) 110675.

[32] A. Muendlein, L. Severgnini, T. Decker, C. Heinzle, A. Leiherer, K. Geiger, H. Drexel, T. Winder, P. Reimann, F. Mayer, C. Nonnenbroich, T. Dechow, Circulating syndecan-1 and glypican-4 predict 12-month survival in metastatic colorectal cancer patients, Front Oncol 12 (2022) 1045995.

[33] S.R. Saroja, K. Gorbachev, T. Julia, A.M. Goate, A.C. Pereira, Astrocyte-secreted glypican-4 drives APOE4-dependent tau hyperphosphorylation, Proc Natl Acad Sci U S A 119(34) (2022) e2108870119.

[34] L. Tatenhorst, F. Maass, H. Paul, V. Dambeck, M. Bahr, R. Dono, P. Lingor, Glypican-4 serum levels are associated with cognitive dysfunction and vascular risk factors in Parkinson’s disease, Sci Rep 14(1) (2024) 5005.

[35] Y. Tamori, M. Kasuga, Glypican-4 is a new comer of adipokines working as insulin sensitizer, J Diabetes Investig 4(3) (2013) 250–1.

[36] H.J. Yoo, S.Y. Hwang, G.J. Cho, H.C. Hong, H.Y. Choi, T.G. Hwang, S.M. Kim, M. Bluher, B.S. Youn, S.H. Baik, K.M. Choi, Association of glypican-4 with body fat distribution, insulin resistance, and nonalcoholic fatty liver disease, J Clin Endocrinol Metab 98(7) (2013) 2897–901.

[37] H. Kaplon, J.M. Reichert, Antibodies to watch in 2019, MAbs 11(2) (2019) 219–238.

[38] C. Hamers-Casterman, T. Atarhouch, S. Muyldermans, G. Robinson, C. Hamers, E.B. Songa, N. Bendahman, R. Hamers, Naturally occurring antibodies devoid of light chains, Nature 363(6428) (1993) 446–8.

[39] F. Khodabakhsh, M. Behdani, A. Rami, F. Kazemi-Lomedasht, Single-Domain Antibodies or Nanobodies: A Class of Next-Generation Antibodies, Int Rev Immunol 37(6) (2018) 316–322.

[40] S. Muyldermans, Nanobodies: natural single-domain antibodies, Annu Rev Biochem 82 (2013) 775–97.

[41] E. De Genst, K. Silence, K. Decanniere, K. Conrath, R. Loris, J. Kinne, S. Muyldermans, L. Wyns, Molecular basis for the preferential cleft recognition by dromedary heavy-chain antibodies, Proc Natl Acad Sci U S A 103(12) (2006) 4586–91.

[42] D.I. Frecot, T. Froehlich, U. Rothbauer, 30 years of nanobodies - an ongoing success story of small binders in biological research, J Cell Sci 136(21) (2023).

[43] M. Feng, W. Gao, R. Wang, W. Chen, Y.G. Man, W.D. Figg, X.W. Wang, D.S. Dimitrov, M. Ho, Therapeutically targeting glypican-3 via a conformation-specific single-domain antibody in hepatocellular carcinoma, Proc Natl Acad Sci U S A 110(12) (2013) E1083–91.

[44] J. Kim, A. Magli, S.S.K. Chan, V.K.P. Oliveira, J. Wu, R. Darabi, M. Kyba, R.C.R. Perlingeiro, Expansion and Purification Are Critical for the Therapeutic Application of Pluripotent Stem Cell-Derived Myogenic Progenitors, Stem Cell Reports 9(1) (2017) 12–22.

[45] G. Behar, P. Chames, I. Teulon, A. Cornillon, F. Alshoukr, F. Roquet, M. Pugniere, J.L. Teillaud, A. Gruaz-Guyon, A. Pelegrin, D. Baty, Llama single-domain antibodies directed against nonconventional epitopes of tumor-associated carcinoembryonic antigen absent from nonspecific cross-reacting antigen, FEBS J 276(14) (2009) 3881–93.

[46] K. Even-Desrumeaux, D. Nevoltris, M.N. Lavaut, K. Alim, J.P. Borg, S. Audebert, B. Kerfelec, D. Baty, P. Chames, Masked selection: a straightforward and flexible approach for the selection of binders against specific epitopes and differentially expressed proteins by phage display, Mol Cell Proteomics 13(2) (2014) 653–65.

[47] J. Schindelin, I. Arganda-Carreras, E. Frise, V. Kaynig, M. Longair, T. Pietzsch, S. Preibisch, C. Rueden, S. Saalfeld, B. Schmid, J.Y. Tinevez, D.J. White, V. Hartenstein, K. Eliceiri, P. Tomancak, A. Cardona, Fiji: an open-source platform for biological-image analysis, Nat Methods 9(7) (2012) 676–82.

[48] J. Su, Y. Song, Z. Zhu, X. Huang, J. Fan, J. Qiao, F. Mao, Cell-cell communication: new insights and clinical implications, Signal Transduct Target Ther 9(1) (2024) 196.

[49] J.P. Buhl, A. Garten, J. Kratzsch, W. Kiess, M. Penke, How Reliable are Commercially Available Glypican4 ELISA Kits?, Exp Clin Endocrinol Diabetes 130(2) (2022) 110–114.

[50] M. Schuksz, M.M. Fuster, J.R. Brown, B.E. Crawford, D.P. Ditto, R. Lawrence, C.A. Glass, L. Wang, Y. Tor, J.D. Esko, Surfen, a small molecule antagonist of heparan sulfate, Proc Natl Acad Sci U S A 105(35) (2008) 13075–80.

[51] I. Farhy-Tselnicker, A.C.M. van Casteren, A. Lee, V.T. Chang, A.R. Aricescu, N.J. Allen, Astrocyte-Secreted Glypican 4 Regulates Release of Neuronal Pentraxin 1 from Axons to Induce Functional Synapse Formation, Neuron 96(2) (2017) 428–445 e13.

[52] K. Ma, S. Xing, Y. Luan, C. Zhang, Y. Liu, Y. Fei, Z. Zhang, Y. Liu, X. Chen, Glypican 4 Regulates Abeta Internalization in Neural Stem Cells Partly via Low-Density Lipoprotein Receptor-Related Protein 1, Front Cell Neurosci 15 (2021) 732429.

[53] I. Jovcevska, S. Muyldermans, The Therapeutic Potential of Nanobodies, BioDrugs 34(1) (2020) 11–26.

[54] R. Kontermann, S. Dübel, Antibody Engineering Volume 2 Springer Berlin, Berlin, 2014.

